# Including biotic interactions in species distribution models improves the understanding of species niche: a case of study with the brown bear in Europe

**DOI:** 10.1101/2023.03.10.532098

**Authors:** Pablo M. Lucas, Wilfried Thuiller, Matthew V. Talluto, Ester Polaina, Jörg Albrecht, Nuria Selva, Marta De Barba, Luigi Maiorano, Vincenzo Penteriani, Maya Guéguen, Niko Balkenhol, Trishna Dutta, Ancuta Fedorca, Shane C. Frank, Andreas Zedrosser, Ivan Afonso-Jordana, Hüseyin Ambarlı, Fernando Ballesteros, Andriy-Taras Bashta, Cemal Can Bilgin, Neda Bogdanović, Edgars Bojārs, Katarzyna Bojarska, Natalia Bragalanti, Henrik Brøseth, Mark W. Chynoweth, Duško Ćirović, Paolo Ciucci, Andrea Corradini, Daniele De Angelis, Miguel de Gabriel Hernando, Csaba Domokos, Aleksander Dutsov, Alper Ertürk, Stefano Filacorda, Lorenzo Frangini, Claudio Groff, Samuli Heikkinen, Bledi Hoxha, Djuro Huber, Otso Huitu, Georgeta Ionescu, Ovidiu Ionescu, Klemen Jerina, Ramon Jurj, Alexandros A. Karamanlidis, Jonas Kindberg, Ilpo Kojola, José Vicente López-Bao, Peep Männil, Dime Melovski, Yorgos Mertzanis, Paolo Molinari, Anja Molinari-Jobin, Andrea Mustoni, Javier Naves, Sergey Ogurtsov, Deniz Özüt, Santiago Palazón, Luca Pedrotti, Aleksandar Perović, Vladimir N. Piminov, Ioan-Mihai Pop, Marius Popa, Maria Psaralexi, Pierre-Yves Quenette, Georg Rauer, Slaven Reljic, Eloy Revilla, Urmas Saarma, Alexander P. Saveljev, Ali Onur Sayar, Cagan H. Şekercioğlu, Agnieszka Sergiel, George Sîrbu, Tomaž Skrbinšek, Michaela Skuban, Anil Soyumert, Aleksandar Stojanov, Egle Tammeleht, Konstantin Tirronen, Aleksandër Trajçe, Igor Trbojević, Tijana Trbojević, Filip Zięba, Diana Zlatanova, Tomasz Zwijacz-Kozica, Laura J. Pollock

## Abstract

Biotic interactions are expected to influence species’ responses to climate change, but they are usually not included when predicting future range shifts. We assessed the importance of biotic interactions to understand future consequences of climate and land use change for biodiversity using as a model system the brown bear (*Ursus arctos*) in Europe. By including biotic interactions using the spatial variation of energy contribution and habitat models of each food species, we showed that the use of biotic factors considerably improves our understanding of the distribution of brown bears. Predicted future range shifts, which included changes in the distribution of food species, varied greatly when considering various scenarios of change in biotic factors, warning about future indirect climate change effects. Our study confirmed that advancing our understanding of ecological networks of species interactions will improve future scenarios of biodiversity change, which is key for conserving biodiversity and ecosystem services.

## INTRODUCTION

In the current biodiversity crisis^1, 2^, understanding how the distribution of species will be impacted by global changes, such as climate and land use changes^3, 4^, is critical for conserving biodiversity and securing associated ecosystem services^5–7^. One of the major outstanding challenges is to capture the complexity of biological responses when making predictions about how species will respond to global changes. Species distributions are indeed governed by a complex set of abiotic and biotic factors^8, 9^. However, most existing models predicting how species will respond to global changes are largely based on abiotic factors, especially climate. This is, in part, a result of the perceived differences in scale with abiotic factors thought to operate at larger spatial scales^10, 11^ and biotic factors, such as species interactions, being considered more important at local scales within communities^10, 11^. Such reliance on abiotic factors when explaining large scale species distributions has also resulted from extremely sparse data on species interactions. This knowledge gap on species interactions, also termed the Eltonian shortfall (understood as the lack of knowledge on intra- and interspecific interactions, but also as the physiological tolerances of species and the effects of species on ecosystems), severely limits our understanding of large-scale biodiversity patterns^12^.

Despite these limitations, studies are beginning to incorporate species interactions into species distribution models (SDMs), which are the most widely used tool for understanding the role of environmental factors in the geographic distribution of species and for predicting potential shifts due to global changes^13^. These studies have demonstrated that adding information on other species into SDMs adds explanatory and predictive power to models^14–16^, suggesting that species interactions are a valuable component in predictive models^17^. However, the approaches used to include species interactions into SDMs usually face three important limitations: (1) they assume spatial co-occurrence among species as interaction^18, 19^, (2) they typically use a binary measure for interaction, e.g., there is/there is not interaction between *Species_A_* and *Species_B_*, and (3) it is assumed there is no spatial variation in the interaction, e.g., interaction between *Species_A_* and *Species_B_* is considered constant in all ecosystems^20^. These latter three processes would be more accurately quantified on the basis of: (1) real data on ecological interactions, e.g., studying the trophic interactions among species; (2) describing the interactions among species with quantitative measures, e.g., measuring the relative energy obtained from food/consumed species; and (3) incorporating the spatial variability of those interactions among different ecosystems^21, 22^, e.g., measuring different values of the relative energy from food/consumed species across geographic space. Therefore, to advance our understanding of species distributions, it is necessary to explicitly account for both abiotic and biotic factors^23, 24^ based on detailed knowledge (Fig. 1).

**Fig. 1.**
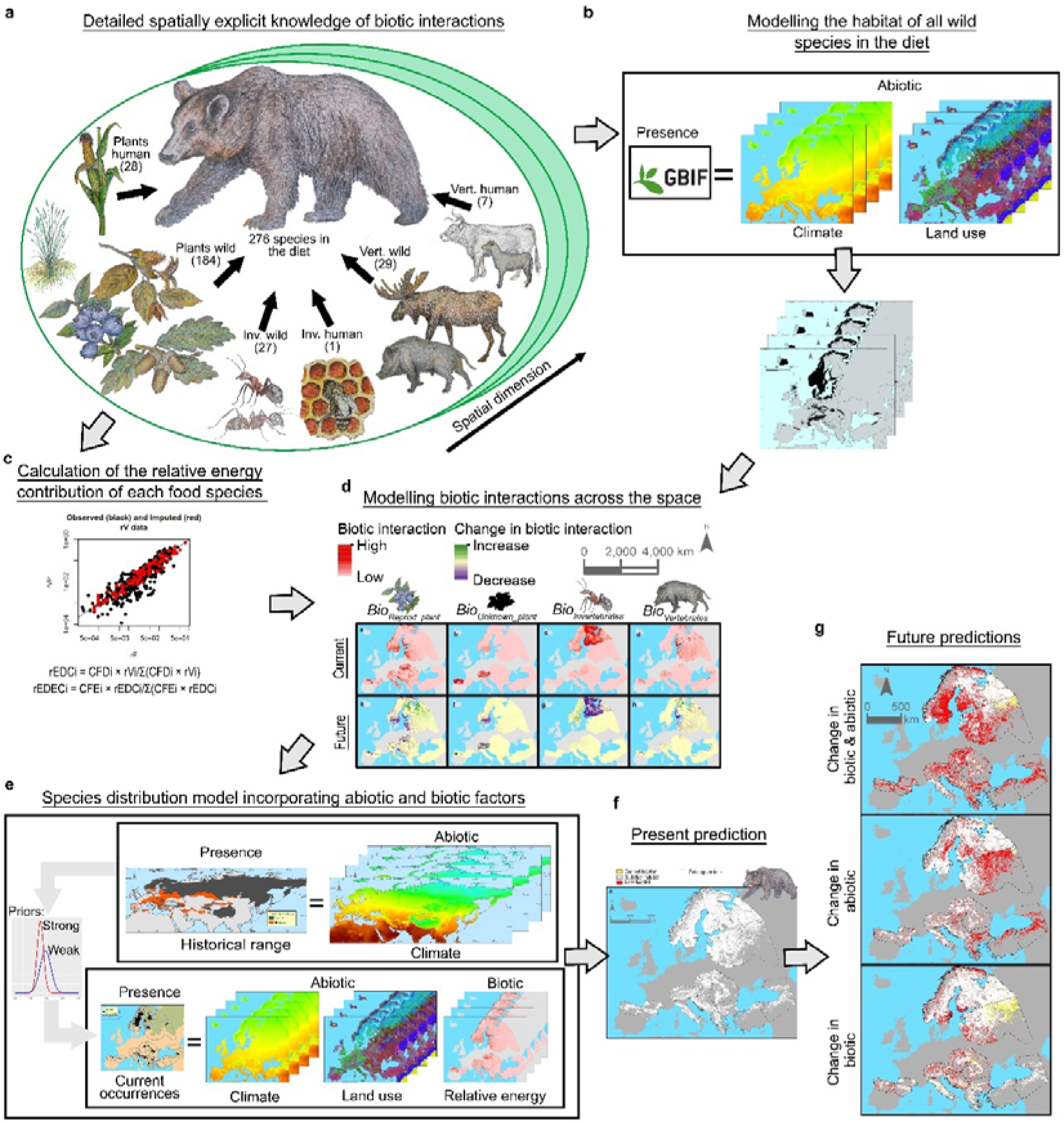
Diagram showing our model system to assess the importance of biotic interactions in understanding the consequences of global change for biodiversity. a, Construction of a database with detailed explicit knowledge of biotic interactions (in our model system brown bear food species in Europe) based on a literature review which accounts for the spatial variability of interactions. b, Fit ensemble SDMs for wild food species and calculation of habitat suitability for the current and three future (2040) Shared Socioeconomic Pathways (SSPs), which were included using predictions for climate^29^ and land use changes^30^. c, Calculation of the relative energy contribution of each food species in different subpopulations/space. d, Calculations of quantitative and binary proxies of biotic interactions across the space and predicting the spatial biotic interactions for each scenario. e, Fit of a species distribution model for the brown bear combining historical and current data and incorporating abiotic factors, which refer to the effects of global changes that directly impact the brown bear including temperature changes (i.e., affecting hibernation and reproduction) and land use changes (i.e., decreasing suitable habitat), and biotic factors, which refer to the effects of global changes through biotic interactions such as changes in the availability of other species as food sources. f and g, Current and future predictions for brown bear distribution considering both abiotic and biotic factors.

Here, we assess whether considering real data on biotic interactions at large spatial scales helps to understand future consequences of global change for biodiversity. Specifically, we tested (i) how biotic interactions change over space, (ii) whether species’ geographic distributions are better explained by quantitative or binary proxies of biotic interactions, (iii) whether species geographic distributions are better explained when combining biotic and abiotic factors, and (iv) whether or not future range shifts differ when considering biotic factors.

We assess these questions using trophic interactions as they are among the most important in determining species distributions and are fundamental to ecosystems^25, 26^. We use the brown bear *Ursus arctos* in Europe as our model system for several reasons: it is a very well-studied species/area, from an ecosystem perspective the brown bear is a top predator and an omnivorous generalist species interacting through its diet with many species, with a very strong impact on ecosystems^27^, and from a conservation point of view, the brown bear is a keystone species in the best conservation areas of the continent^27^ and several of its subpopulations are at risk of extinction^28^. Thus, advances in our model system will target a wide range of species and communities with high conservation value. We constructed a database of more than 3 million high resolution (1 km^2^) brown bear occurrences with data from 14 subpopulations across Europe and Turkey (*Occurrence Database*; Supplementary Tables 1-3 and Supplementary Fig. 1; Methods). To obtain detailed knowledge of biotic interactions, we reviewed 47 studies of brown bear diet in this area, constructed a unique, highly detailed, spatially-explicit database of trophic interactions (*Trophic Database*; Figs. 1a and 2; Supplementary Tables 4-7 and Supplementary Fig. 2), and calculated the relative energy contribution of each food item (Fig. 1c). To address question (i), namely whether biotic interactions change over space and if they are explained by environmental factors, we used averaged linear models predicting the relative energy contribution of different food categories (i.e., reproductive plants, vegetative plants, unknown plants, invertebrates and vertebrates) and the diet diversity of these food categories as a function of climate^29^ and land use^30^ variables. To assess (ii), whether species geographic distributions are better explained by quantitative or binary proxies of biotic interactions, for each wild food species in the *Trophic Database*, we fitted ensemble SDMs at 1 km^2^ using GBIF occurrences^31, 32^ and climate and land use variables^29, 30^ as predictors, and calculated the habitat suitability for the current and three future Socioeconomic Shared Pathways (SSP; SSP1-2.6, SSP3-6.0 and SSP5-8.5), considering climate and land use change (Fig. 1b; Supplementary Figs. 3-5). We calculated two measures of biotic interactions. The first was a quantitative measure of biotic interactions (*Biotic variables*) obtained by multiplying the relative energy contribution described at the species level in each subpopulation (defined as parts of the distribution of the species that are isolated from others and/or present different environmental characteristics and/or conservation status; Fig. 2a; See Supplementary Methods and Supplementary Fig. 2) by the current habitat suitability of each species, and then combining the values for each food category (Fig. 2d). The second was a binary measure of biotic interactions (*Biotic_binary)* calculated by multiplying the current habitat suitability by 1 or 0 depending on, respectively, whether or not an interaction with food species was observed in each subpopulation, and then combining the values for each food category. For each food category, we fitted and evaluated (AIC based) two univariable SDMs explaining brown bear distribution using the brown bear *Occurrence Database*: (i) a model using *Biotic variables* as predictors and (ii) a model using *Biotic_binary* variables as predictors. To assess (iii), whether species’ geographic distributions are better explained when combining biotic and abiotic factors, we fitted and evaluated (WAIC based) three Bayesian models (BMs) to explain brown bear distribution, using data from the brown bear *Occurrence Database* as a response variable: (1) a model with abiotic (climate and land use variables) and biotic predictors (using the best overall proxies, *Biotic variables* or *Biotic_binary*, from the previous univariable SDMs), (2) a model with abiotic predictors only, and (3) a model with biotic predictors only (Fig. 1e). To minimize bias or the truncation of the environmental space when using only current data^33–35^, two of these BMs utilized historical range data (*Range Database*): the abiotic and biotic BM, and the abiotic BM. Historical range data was included using a Bayesian hierarchical model (BHM) which combined models with historical geographic range as a response variable and historical climate variables as predictors and models with current data. To address (iv), whether future range shifts differ when considering biotic factors or not, we used the best BMs and assessed changes in the potential distribution of the brown bear combining the three previous SSPs with three scenarios of change: (1) change in abiotic and biotic variables, (2) change in abiotic variables and (3) change in biotic variables (Fig. 1f and 1g; Methods).

**Fig. 2.**
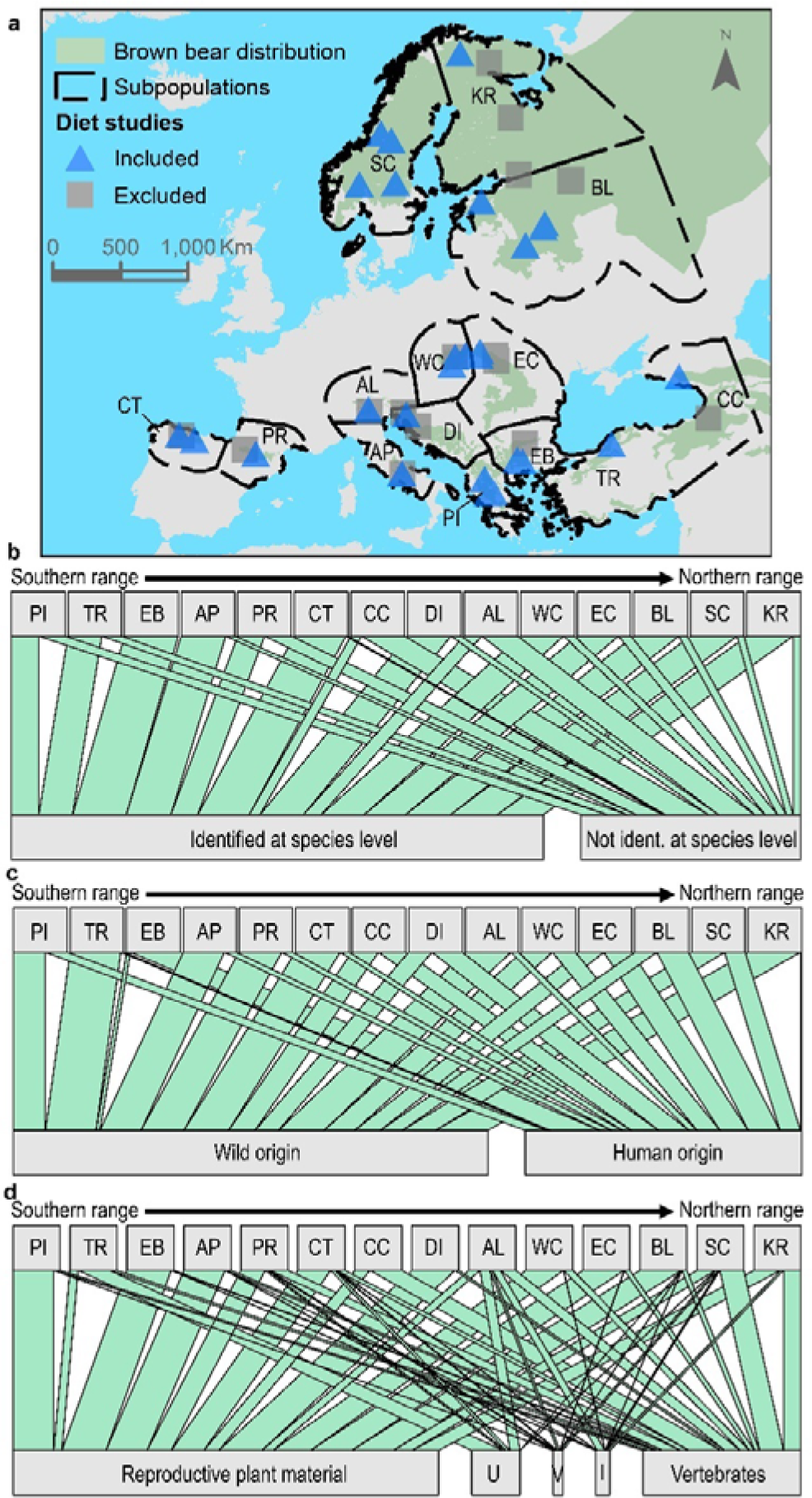
Brown bear food web in Europe. a. Map showing the 14 brown bear subpopulations considered: Pindus (PI), Turkey (TR), East Balkan (EB), Apennine (AP), Pyrenees (PR), Cantabrian (CT), Caucasian (CC), Dinaric (DI), Alpine (AL), Western Carpathian (WC), Eastern Carpathian (EC), Baltic (BL), Scandinavian (SC) and Karelian (KR). We also mark the location of all the studies of brown bear diet reviewed, indicating if they were ultimately included (n = 31) or not (n = 16); note that not all studies are visible due to the overlapping of locations. **b,** Relative estimated dietary energy content (*rEDEC*, a proxy for the relative importance of each species) identified or not identified at the species level for each of the 14 brown bear subpopulations in Europe. **c,** Proportion of the *rEDEC* for wild species and those of human origin. **d,** Proportion of the *rEDEC* identified at the species level for each food category (U for unknown plant material, V for vegetative plant material, I for invertebrates), for each of the 14 brown bear subpopulations in Europe.

**Fig. 3.**
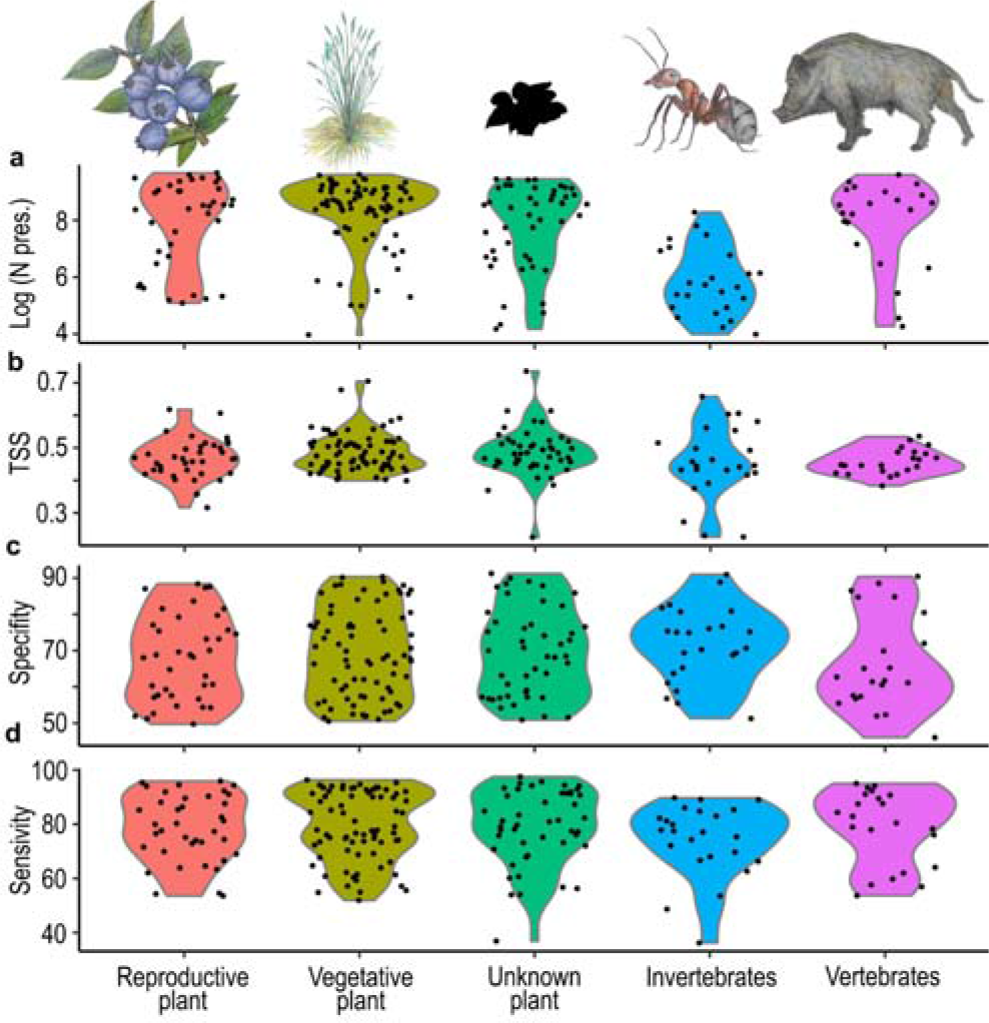
Number of presences and validation of SDMs for wild food species according to category (i.e., reproductive plants, vegetative plants, unknown plant material, invertebrates and vertebrates). a, Number of presences from the GBIF used to fit a species distribution model (SDM) for each of the 205 wild food species out of the total 240 wild species recorded (85.4% of the wild species in the brown bear diet with enough data to fit a SDM). b, Predictive quality of SDMs using the true statistics skill (TSS), c, Specificity (percentage of absences correctly predicted) and d, Sensitivity (percentage of presences correctly predicted).

## RESULTS

### Spatial variation of biotic interactions

From the literature search, we identified trophic interactions between the brown bear and 276 food species (Fig. 1; Supplementary Table 8). In total, 76.1% of species were plants and 23.2% were animals (13.0% vertebrates, 10.1% invertebrates). When focusing only on trophic interactions described at the species level, we found that the relative energy contribution of each food category varied among subpopulations; e.g., in the Scandinavian subpopulation, 51% of the energy was of vertebrate origin compared to just 4% for the East Balkan subpopulation (Supplementary Tables 9 - 11). Also, the proportion of energy from human-derived sources (n = 36 species) strongly varied among subpopulations; e.g., in the Karelian subpopulation only 2% of the energy was from human-derived sources compared to 93% for the East Balkan subpopulation (Fig. 2a-2c; Supplementary Table 12).

Using all food items in each study site (n > 1,300; not only those described at the species level), we found that the relative energy contribution to the diet of the brown bear for all food categories (apart from the vegetative plant category), as well as the diversity of food categories, was driven by climate and land use (climate and land use variables showed p-values < 0.05; Supplementary Tables 13-32). For example, bears consumed proportionally more invertebrates and fewer reproductive plant parts (e.g., fleshy fruits and nuts) in areas with low vegetation cover, and proportionally more vertebrates in areas with climates exhibiting a smaller diurnal temperature range and in areas with more shrubland and more sparce vegetation. Similarly, bears tend to have more diverse diets when they occur in areas with both high broadleaved forest cover (broadleaved forest showed p-values < 0.05 in the three models explaining each diversity index; Supplementary Tables 30-32) and a wide diurnal temperature range (diurnal temperature range showed p-values < 0.05 in the models explaining diversity indexes; Supplementary Tables 30-31).

### Quantitative versus binary proxies of biotic interactions to explain geographic distributions

Among all wild food species (n = 240; Supplementary Table 8), we were able to build robust SDMs and predict the current/future habitat suitability for 205 species (*SDM_Food_*: average sensitivity of 78.4%, specificity of 69.2% and TSS of 0.48). Invertebrate species were, however, underrepresented in the GBIF data, which affected the quality of the predictions for these species (Fig. 3; Supplementary Tables 33-39). However, since they were of minor importance in the diet of the brown bear in Europe, with an average of only 2% among all subpopulations (Fig. 2d and Supplementary Table 11), we believe that this does not affect our main conclusions. We then contrasted whether brown bear distribution was better explained by quantitative (*Biotic variables*) or binary proxies of biotic interactions (*Biotic_binary*). For the vegetative plant food category, we found that the binary proxy for the interaction was sufficient to best explain brown bear distribution. Conversely, for all other food categories (i.e., reproductive plants, unknown plants, invertebrates and vertebrates), including a quantitative measure of the interactions with species better explained the distribution of brown bears (Supplementary Table 40).

### The role of abiotic and biotic factors in explaining species’ geographic distributions

When we compared the three Bayesian models (BMs), we found that the model combining abiotic and biotic factors was the best and significantly improved the understanding of brown bear distribution compared to models using either abiotic or biotic factors (delta WAIC*_Abiotic_*= 337.1; delta WAIC*_Biotic_* = 1,611.2; Supplementary Tables 42-53 and Supplementary Figs. 7-12). The model combining abiotic and biotic factors showed a good performance compared with a *Null* model using only the intercept (delta WAIC*_Null_* = 2,539.7) and yielded a good classification accuracy (0.56; Supplementary Table 54), with a low rate to correctly classify the pseudo-absences of brown bear (true negative rate = 0.25) and a high rate to correctly classify the presences of brown bear (true positive rate = 0.87). The model/threshold for classifying presence/absence was very good at predicting the presence of species, but tended to overestimate it from its current distribution, which was intentional, and similarly to other studies of top predators which have suffered important range contraction^36^, as the past direct persecution of the species has locally eliminated the population from potentially suitable areas^34^ (see Methods). The predictions showed a current potential distribution for the brown bear of 2,793,351 km^2^ (Fig. 4), of which 68% overlaps with the known area of occupancy. Important differences for each subpopulation were noted, e.g., the Karelian habitat is 98% occupied, but the Alpine, Cantabrian, Apennines and Pyrenees habitats are only 8.9%, 14.5%, 8.7% and 15.8% occupied, respectively. In other words, there are large areas in those mountain regions that could host brown bears but currently do not. Importantly, for a species with several subpopulations at risk of extinction, only 19% of the potential distribution is within protected areas.

**Fig. 4.**
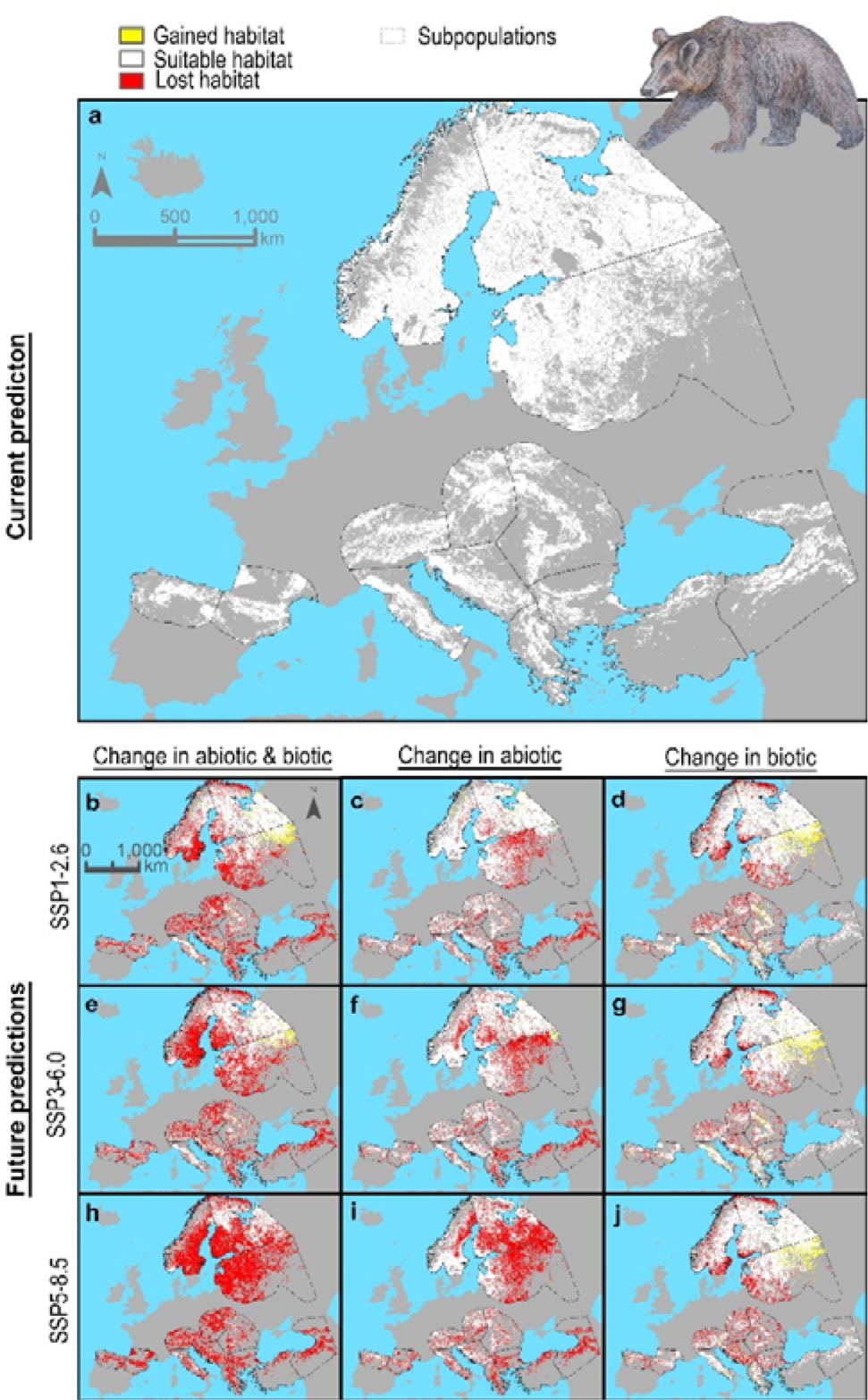
**Brown bear habitat predictions. a**, Prediction of brown bear habitat for current conditions. Future predictions of habitat for climate change scenarios SSP1-2.6, SSP3-6.0 and SSP5-8.5 considering changes in both abiotic and biotic factors (**b**, **e**, **h**), changes in abiotic factors only (**c**, **f**, **i**) and changes in biotic factors only (**d**, **g**, **j**). The predicted area only includes a buffer area of 200 km around the current distribution to avoid extrapolating biotic variables into an environmental space where there is no information about the trophic interactions of brown bears.

In terms of species response to the selected abiotic and biotic variables, brown bear presence showed a bell-shaped response to most climate variables, but as expected, a negative response to the percentage of urban areas, and a positive association with forests and natural landscapes (Fig. 5). While the response of brown bear presence to most biotic variables/food categories was positive (i.e., reproductive plants, unknown plants and vertebrates), it showed a surprising negative association with the invertebrate food category (see Discussion and Supplementary Table 49).

**Fig. 5.**
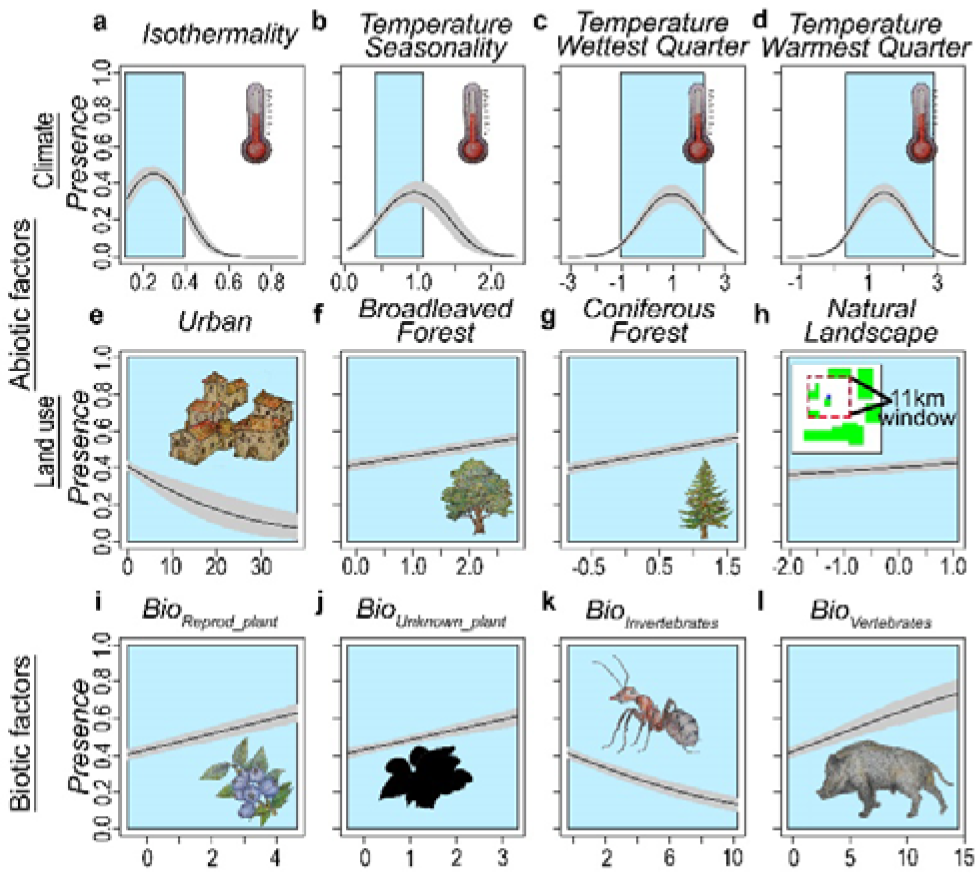
Partial response plots of brown bear distribution to both abiotic and biotic variables. The distribution model for brown bear including both abiotic and biotic factors was fitted combining both historical (*Range Database*) and current data (*Occurrence Database*). The continuous line represents the mean response value and the grey area shows the model uncertainty (95% confidence interval). The blue area indicates the range of values of the current data. Isothermality (*Clim_3*), temperature seasonality (*Clim_4*), mean temperature of the wettest quarter (*Clim_5*) and mean temperature of the warmest quarter (*Clim_10*).

### The effect of biotic factors in future range shifts

The predictions of biotic interactions, based on the best measures of interaction (*Biotic variables*), for future SSPs showed important differences. The potential available energy for the brown bear from all food species (*Bio_All_species_)* was predicted to be reduced by 53% under SSP3-6.0. Importantly, when focusing on the different food categories, future predictions for biotic variables showed differences by food category and spatially varied by subpopulation e.g., *Bio_Vertebrates_* for SSP3-6.0 is predicted to be reduced by 83% for the Western Carpathian subpopulation, whereas for the Apennines subpopulation it is predicted to increase by 16% (Fig. 1d; Supplementary Fig. 6 and Supplementary Table 41).

Those predicted future changes in biotic interactions affected the projected distribution of brown bear in the future. Our model showed a drastic range reduction that was more marked when considering both abiotic and biotic variables, with an overall reduction of 50%, as compared to either biotic (reduction of 13%) or abiotic (reduction of 31%) variables only. Range reduction was most pronounced in the south-eastern subpopulations, e.g., 95% reduction in the East Balkan subpopulation and 99% reduction in the Turkey subpopulation (Fig. 4; Supplementary Table 58). Importantly, the spatial variability of future changes in biotic interactions and by food category described above, translated into different effects across the brown bear range, as abiotic and biotic factors acted differently among the different subpopulations. For example, according to the SSP3-6.0 scenario, in the Alpine subpopulation, biotic variables were associated with a habitat reduction of 37% compared to 23% for abiotic variables, whereas in Turkey, biotic variables explained a comparatively smaller habitat reduction of 8% (Supplementary Tables 55-58).

## DISCUSSION

Our analyses demonstrate the importance of biotic interactions in explaining species distributions at continental scales. Specifically, we found that 1) biotic interactions are highly variable across geographic space and are determined by climate and land use variables, 2) reliably estimating biotic factors for SDMs requires accounting for quantitative measures of biotic interactions, 3) including biotic interactions in SDMs significantly improves our understanding of species distributions and 4) the consideration of biotic interactions in future projections has important effects on predicting future consequences of climate and land use changes for species distributions. We also found that both plant- and animal-based interactions are important, demonstrating the need to include both for omnivorous species, and we confirmed the importance of the brown bear as a keystone species in ecosystems.

Climate and human land use determined the biotic interactions through different ecosystems, suggesting that future scenarios of climate and land use may directly affect the interactions of brown bears with food items, having important implications for ecosystem functioning and services, e.g., seed dispersal, reducing the abundance of ungulates or medium-size predators, which can influence the diversity of ecosystems^37, 38^ and/or increase human-wildlife conflicts^39^, which is a problem for the conservation of species^40^.

The distribution of brown bears in Europe depends on climate variables influencing the physiological range of the species^41^, as well as land use, such as forested areas and continuous natural areas which could be used as shelters^27, 36^. However, here we show that even biotic factors shape brown bear distribution, represented here as the relative energy derived from food resources available to the species. According to our results, brown bears select areas that maximise this available energy (i.e., positive response of brown bear to most biotic variables), which may be explained by the high energy requirements of the species^42^ and the absence of strong interspecific competition for food resources^43^. The negative association with the availability of energy from invertebrates may be related to their negative correlation with isothermality (*Clim_3*; Correlation = -0.47), which has great importance for brown bear distribution and exhibits a bell-shaped response (Fig. 5). This could indicate that areas with high available energy from invertebrate species are located in less suitable environmental spaces respect to this abiotic factor. In addition, the low relative energy which invertebrates represent in the brown bear diet —an average of 2% among all subpopulations (Fig. 2d and Supplementary Table 11)— may suggest that they represent opportunistic consumption more than intentional/preferred prey, which may not influence the distribution of brown bears. Our results showed that brown bears currently have a large amount of suitable habitat that could be occupied, especially in southern subpopulations, e.g., Cantabrian and Apennine subpopulations. However, climate and land use changes might cause an important reduction in their suitable habitat, and thus the protection of forests, the reduction of landscape fragmentation and the conservation of the community of species interacting with brown bears represent crucial buffers against these drivers.

The use of SDMs is one of the most advanced and widely used tools to understand the factors delimiting species distributions and to predict the effects of global change on biodiversity^13^. Our results show the importance of additionally considering biotic factors and taking an ecosystem approach to properly understand species distributions^44, 45^, especially for modelling species distributions under climate and land use change scenarios^46^. However, as biotic interactions are highly complex, their inclusion needs to be accounted for on the basis of ecological studies which include their spatial heterogeneity and quantitative estimates^20, 47^. Our approximation was a simplification of the true biotic interaction networks of the brown bear, and other aspects such as competition, parasitism, the potential plasticity of different subpopulations to alter their diet in future scenarios and/or other dimensions including temporal variation at seasonal, interannual and other spatial scales may be relevant and could be considered in future studies. In general, implementation of these new models in other species currently faces two big challenges: (1) detailed and extended information about ecological interaction networks, and (2) high-quality data about species presences/occurrences. To overcome these challenges, global-scale monitoring initiatives with open-source principles, open-source databases on species ecology and the reduction of spatial and taxonomic biases will be of primary importance^48, 49^. This new generation of SDMs, which allow decoupling abiotic and biotic factors, will better identify the drivers responsible for species distributions and will enhance the predictions regarding the effects of global change on species; overall they will generate important knowledge to conserve biodiversity and ecosystem services.

## METHODS

### Biotic interactions

#### Review of brown bear diet studies

We reviewed 47 studies of brown bear diet in Europe by searching in SCI Journals, master’s and PhD theses, and grey literature (research that is either unpublished or has been published in non-commercial form, e.g., technical reports, conference proceedings; Supplementary Tables 4 and 5 and Supplementary Figure 2). For consistency with climate and land use data, we selected 31 studies conducted between 1989 and 2018 which had sufficient taxonomic resolution (genus and/or species). For each study, we recorded three types of information: study area location; the type of samples, e.g., brown bear scat or the stomachs of dead individuals; and the number of samples. Additionally, within each study and for each food item we recorded two parameters: (1) the relative frequency of occurrence (*rF*), calculated as the number of occurrences of food item *i* divided by the total number of occurrences of all food items, that is *rF_i_* = *f_i_*/Σ*f_i_*; and (2) the relative volume (*rV*), calculated as the volume of food item *i* divided by the total volume of all food items, that is *rV_i_*= *v_i_*/Σ*v_i_*.

#### Calculating energy available from food items

Because not all studies reported estimates of *rV*, we used the strong relationship between *rF* and *rV* (r = 0.86, 0.81-0.90 95% CI from bootstrapped correlation coefficients; Supplementary Figures 3-5) to impute *rV* for those studies with missing data. We modelled the relationship between *rV* and *rF* based on studies in which both variables were available with a Bayesian hierarchical model that was fitted using the R package MCMCglmm^50^. We modelled the log(*rV_ij_*) of food item *i* in study *j* (so one prey item can have multiple entries if it was found in multiple studies) using a normal distribution. We modelled log(*rV_ij_*) as a function of log(*rF_ij_*), diet category (i.e., vertebrate, invertebrate, seeds, fruits, vegetation or other [i.e., unidentified material or garbage]), and their interaction. We included a random intercept for study ID. We placed uninformative normal priors on the coefficients of the fixed explanatory variables (i.e., [0, 10^10^]), and weakly informative inverse-gamma priors with shape = rate = 0.001 on the variance components (i.e., IG[0.001, 0.001]). The model was run for 130,000 iterations with a burn-in of 30,000 and a thinning interval of 100 iterations, resulting in 1,000 posterior samples. Convergence was checked using the potential scale reduction factor (PSRF < 1.1)^51^ and temporal autocorrelation (r < 0.1) using the R package coda^52^. We used this model to impute *rV* for those studies with missing data. Then we applied two sets of correction factors to the estimates of *rV* to account for differences in digestibility and energy content between food items. We first applied correction factors for digestibility (*CF_D_*; Supplementary Table 6) to calculate the relative dry weight of each food item *i* in each study (*rEDC*)^53^ using the formula: *rEDC_i_* = *CF_Di_* × *rV_i_*/Σ(*CF_Di_* × *rV_i_*). Then we used a second set of correction factors (*CF_E_*) to convert dry matter to digestible energy, and calculated the relative estimated dietary energy content (*rEDEC*)^54^ of each food item *i* in each study: *rEDEC_i_* = *CF_Ei_* × *rEDC_i_*/Σ(*CF_Ei_* × *rEDC_i_*).

In diet studies, *rVi* and/or *rF_i_* are often provided for groups of several species, and within these groups the species are usually described as present but without quantification of *rF* and/or *rV*, and thus the *rEDEC_i_* provides a description for a broad taxonomic group. To improve the description at the species level, we followed a similar approach to previous food-web studies^55, 56^. For each group containing several species, we assigned its described *rEDEC_i_* to the species (*s*) that was most frequent in the bear diet, *rEDEC_s_ = rEDEC_i_*, when explicitly stated in the paper. If several species belonging to the group were consumed by bears, but none explicitly mentioned as the most frequently eaten or occurring in scat, we divided equally the *rEDEC_i_* among the species consumed by bears and present in the area, *rEDEC_s_ = rEDEC_i,_/n_species_*. For example, in Naves et al. (2006), we equally assigned *rEDEC_Quercus_*among the three *Quercus* species present and reported in the article as being part of the diet, *rEDEC_s_ =* 22.07/3, thus each one was assigned 7.36%. We applied this method to items grouped at the genus level for plants and animals when the result was *rEDEC_s_* > 3% (Supplementary Tables 5 and 7).

#### Calculating associations between diet and environmental variables

We used the selected studies of brown bear diet (Supplementary Table 4) to calculate the association between latitude, land cover and climate variables and the *rEDEC* in each diet category. We first calculated the value of land use^30^ and climate^29^ variables (Supplementary Tables 59 and 60) within a buffer area of 18 km (an area of 1,018 km^2^) around the site locations of the selected studies. We eliminated highly correlated variables using the variance inflation function (VIF)^57^ in the R package usdm^58^. Then, we calculated all possible linear models explaining the percentage of each diet category using all possible combinations of the remaining uncorrelated variables as predictors, using the package MuMIn^59^ in R. Using the subset of best models (delta AICc < 3) we calculated an average model using the subset option. We also calculated models explaining diet diversity (among the diet categories) as a function of land use and climate variables in the buffer areas. We first calculated three indexes of diversity for the diet categories (Simpson, Shannon and Inverse Simpson)^60, 61^ using the R package vegan^62^. We fitted linear models explaining each diversity index as a function of the uncorrelated variables previously calculated. We calculated, for each index, all possible models using all possible combinations from the uncorrelated variables, and using the subset of best models (delta AICc < 3) we calculated an average model using the subset option.

#### Calculation of a representative diet for each subpopulation

The brown bear is a generalist omnivore species that shows high variation in its diet across its geographic distribution^63^. For example, the diet of bears in Scandinavia has a relatively higher proportion of vertebrates compared to individuals living in southern Europe, where the consumption of vegetation is comparatively much higher^63^. To consider this spatial variation in the brown bear diet, we calculated a representative diet of the brown bear for each *subpopulation* (*Subp*; Supplementary Figure 1). To differentiate brown bear subpopulations in Europe we used the definition of subpopulations provided by the IUCN, i.e., “geographically or otherwise distinct groups in a population between which there is little exchange”. We utilized the current geographic distribution of brown bears^28^. Separated polygons were characterised as subpopulations and continuous polygons showing important differences in climate, habitat and conservation status of different spatial entities were split into subpopulations in order to account for this variability (e.g., Scandinavian, Karelian and Baltic subpopulations)^28, 64^. Our assignment of subpopulations was similar to Chapron *et al.*^65^’s subdivisions for the brown bear, but we included two new entities for Europe. First, we considered *Pindos* a distinct subpopulation given its differences in climate and habitat from the *Dinaric* and *East Balkan* subpopulations. Second, we considered *Western Carpathian* and *Eastern Carpathian* brown bears to represent different subpopulations on the basis of differences in habitat, climate and conservation status^28, 66^. Third, two new subpopulations were considered for Turkey, *Turkey* and *Caucasus*. We assigned to each *subpopulation* a unique diet (*D_S_*) on the basis of the reviewed studies for each subpopulation.

Similarly to Banašek-Richter *et al.* 2004^47^, where consumed biomass was used as a quantitative descriptor for the flow of energy in the food system, we used the *rEDEC* previously calculated, which provides a more realistic version of the flow of energy in the brown bear food-web system. From the diet studies within each *subpopulation, Subp*, we calculated for each food species (*S*), the *rEDEC_SubpS_*, and assumed it to be a representative *rEDEC* in that subpopulation:

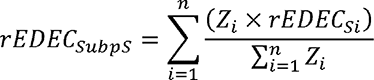

*rEDEC_SubpS_* is the representative *rEDEC* in the subpopulation *Subp* for food species *S*, *i* is each diet study within the subpopulation *Subp* (Supplementary Figure 1 and Supplementary Table 4), *n* is the number of diet studies in the subpopulation *Subp* (Supplementary Table 4), *Z* is the number of sampling units (n scats, or n of stomachs analysed) in each study (we use this term to give more importance to studies with more data; Supplementary Table 4), *rEDEC_Si_*is the *rEDEC* of food species *S* in diet study *i* (Supplementary Table 7).

#### Calculation of habitat suitability for each wild food species

We modelled the habitat of wild food species in the diet database (Supplementary Tables 8 and 33). To do this, we used the R package rgbif^67^ to download occurrences of each food species from the Global Biodiversity Information Facility (GBIF)^31^, a database which is widely used to assess the effects of climate change on biodiversity^68, 69^ for species distribution modelling^70, 71^ and species conservation^72, 73^. We selected all occurrences of food species with coordinates, obtained from human or machine observations (e.g. camera traps), an uncertainty in meters of < 564 m to match the circumference radius of a circle of an area of our cell size (1km^2^), located in Europe, North Africa and the Middle East (between decimal latitude “15, 75” and decimal longitude “-20, 105”) and for the period 1989–2018. Then, we used package CoordinateCleaner^74^ to remove usual errors in occurrences (i.e. country centroids). Points of occurrence were transformed into presences and pseudo-absences to reduce the potential spatial bias^75–77^ by aggregating points into equal-area grid cells of 1x1 km using a Conic Equal Area projection (Europe Albers Equal Area Conic). Each grid cell that contained at least one occurrence point was assigned a ‘1’, and cells without any record of occurrence were assigned a ‘0’. Furthermore, we excluded species with <50 ‘presence’ grid cells from the analyses.

For each of these food species, we selected the same number of random pseudo-absences as presences inside a minimum 3 km-radius plot and a maximum 10 km-radius plot at each presence location. We preselected a total of 11 environmental explanatory variables. First, we preselected a subset of 4 bioclimatic variables from the CHELSA dataset^29^ following previous studies^78^ (Supplementary Table 59), and then we preselected 7 land use/land cover variables representing the percent cover of different land use categories (Supplementary Table 60), obtained from the Globio4 dataset^30^. We excluded land use/land cover representing the percent cover of urban areas to avoid the introduction of a land cover category potentially oversampled.

To build the habitat model for each food species, we selected the 6 best variables among the previous 11 environmental explanatory variables using the following procedure. First, we calculated the variance inflation factor (VIF)^57^ and excluded highly correlated variables (VIF > 10) using a stepwise procedure in the R package usdm^58^. Second, from the remaining uncorrelated variables, we fitted univariate Generalized Linear Models (GLMs) and selected the best 6 variables using an information theoretic approach based on Akaike Information Criterion (AIC)^79^. Using these 6 selected variables, we modelled the habitat of each food species applying ensemble modelling, a statistical technique that improves the robustness of predictions^80^, using the R package Biomod2^81^. For each species we calculated 12 models: (a) we repeated the selection of pseudo-absences twice, each time selecting the same number of random pseudo-absences as presences inside the 3-km and 10-km radius plots (as in the univariate models); (b) for each selection of pseudo-absences we repeated the process of data splitting twice (taking 70% of pseudo-absences to fit the model and 30% to evaluate); and (c) for each dataset of data splitting we fitted three different modelling algorithms: GLM with quadratic and second order polynomials allowed (for all predictors), Generalized Boosting Model/Boosted Regression Trees (GBM) with 3,000 trees and Random Forest (RF) with 750 trees. Each of the 12 fitted models (2 pseudo-absences selection x 2 data splitting x 3 modelling algorithms) was evaluated using the true skill statistic (TSS)^82^ and we selected models with TSS > 0.2 to include in the ensemble-model building.

Using the fitted ensemble models, we predicted the habitat suitability of each food species for the current and three future climate and land use scenarios. Future scenarios were obtained by using the climate data from the Institut Pierre Simon Laplace Model CM5A-MR (IPSL-CM5A-MR)^83^ from the CHELSA^29^ database and land use forecasts from the Globio4^30^ database for the year 2050.

The Globio4^30^ database uses three different scenarios based on a combination of three Shared Socioeconomic Pathways (SSPs), which are scenarios of projected socioeconomic global change up to the year 2100, including (1) the Sustainability scenario, SSP1; (2) the Regional Rivalry scenario, SPP3; and (3) the Fossil-Fuelled Development scenario, SPP5^84, 85^; with three Representative Concentration Pathways (RCPs)^86^: (1) the very stringent scenario, RCP2.6; (2) the intermediate stabilisation pathway scenario, RCP6.0; and (3) the scenario of comparatively high greenhouse gas emissions, RCP8.5^4, 84, 87^. As a result, combined with the Globio4 database has the SPP1_RCP2.6 scenario, the SPP3_RCP7.0 scenario and the SPP5_RCP8.5 scenario.

From the IPSL-CM5A-MR from CHELSA, we selected the RCP2.6 scenario, RCP6.0 scenario and the RCP8.5 scenario. Thus, combining the future scenarios from the Globio4 database and from CHELSA we obtained three future scenarios for land use and climate, namely SSP1-2.6, SSP3-6.0 and SSP5-8.5.

#### Modelling the potential energy available to the brown bear across space

We used two proxies to calculate spatial variables describing the biotic interactions between the brown bear and food species in its diet. The first approximation generated the *Biotic variables*, which were based on detailed quantitative criteria using a combination of *rEDEC_SubpS_* and the habitat suitability prediction. The second approximation generated the *Biotic_binary variables*, which were based on a qualitative measure, a binary link using the presence/absence of interaction between each food species and the brown bear for each subpopulation and the habitat suitability prediction.

#### Calculating potential biotic interactions on the basis of quantitative links: Biotic variables

We used the previously calculated *rEDEC_SubpS_*(See section *Calculation of a representative diet for each subpopulation*) as a link representing the interaction strength between food species *S* and the consumer, the brown bear, among the subpopulation *Subp*.

The spatial distribution of these interactions is heterogeneous across the landscape as it depends on the co-occurrence of the food species and the consumer, and a higher co-occurrence of both species will be positively associated with interaction strength^20^. Thus, within each subpopulation, we assigned the calculated *rEDEC_SubpS_*as a potential link across the predicted habitat of food species *S*, multiplying the *rEDEC_SubpS_* by the habitat suitability of food species *S*. Thus, for each grid cell (*C*) of each *subpopulation* we calculated a term called *Potential energy* (*Pe*) as:

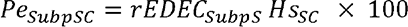

where:

*Pe_SubpSC_* is the potential energy in the subpopulation *Subp* for species *S* in cell *C*,

*Hs_SC_* is the habitat suitability of species *S* in cell *C*.

We considered groups of food species (i.e., reproductive plants, vegetative plants, unknown plants, invertebrates and vertebrates, and a group including all food species) which are vital for the brown bear and calculated the *Sum of the Potential energy* for all food species in each group *G* as:

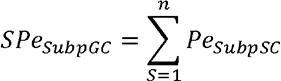

where:

*SPe_SubpGC_* is the sum of the potential energy in the subpopulation *Subp* for all food species in group *G* in cell *C*,

*n* is the number of food species in each subpopulation *Subp* and group *G*, *G* represents each of the groups of food species in the diet considered (reproductive plants, vegetative plants, unknown plants, invertebrates and vertebrates) and a group including all food species.

Using the *SPe_SubpGC_* over all cells of our study area, we obtained the *Biotic variables*, which represent a spatial downscaling of the potential interaction strength among the food species of each diet group considered. Thus, we obtained six *Biotic variables: Bio_All_species_*, *Bio_Reprod_plant_*, *BioVeget_plant*, *BioUnknown_plant*, *BioInvertebrates*, and *BioVertebrates*.

*Calculating potential biotic interactions on the basis of binary links: Biotic_binary variables* We used a categorical/binary description for each subpopulation, *Binary_SubpS_*, as a link representing the interaction strength between food species *S* and the brown bear within the subpopulation *Subp*. *Binary_SubpS_* has the binary value 1 when there was consumption of food species *S* by brown bear in the *Subp*, and 0 when there was no consumption of food species *S* by brown bears in the *Subp*.

In this case, within each *subpopulation*, we assigned the calculated *Binary_SubpS_* as a potential link over the predicted habitat of food species *S*, multiplying the *Binary_SubpS_*by the habitat suitability of food species *S*. Thus, for each grid cell (*C*) of each *subpopulation* we calculated a term called *Potential energy* (*Pe*) as:

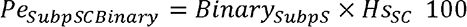

where:

*Pe_SubpSCBinary_* is the potential energy in the subpopulation *Subp* for food species *S* in cell *C*,

*Hs_SC_* is the habitat suitability of food species *S* in cell *C*.

We used the same diet groups of food species considered above for *Biotic variables*, and we calculated the *Sum of the Potential energy Binary* for all food species in each group *G* as:

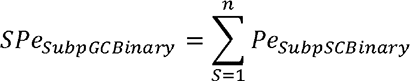

where:

*SPe_SubpGCBinary_* is the sum of the potential energy in the subpopulation *Subp* for all food species in group *G* in cell *C,*

*n* is the number of food species in each subpopulation *Subp* and group *G*,

*G* represents each of the groups of food species in the diet considered (reproductive plants, vegetative plants, unknown plants, invertebrates and vertebrates) and a group including all species.

Using the *SPe_SubpGCBinary_* over all cells of our study area we obtained the *Biotic_binary variables*, which represents a spatial downscaling of the potential interaction strength among the food species in each diet group considered. Thus, we obtained six *Biotic_binary variables: BioB_All_species*, *BioB_Reprod_plant*, *BioB_Veget_plant*, *BioB_Unknown_plant*, *BioB_Invertebrates*, and *BioB_Vertebrates*.

#### Comparison of biotic proxies explaining brown bear distribution

To assess which proxies of biotic interactions better explained brown bear distribution, we compared for each group *G*, the univariate models for *Biotic variables and Biotic_binary variables*. Models were fitted using a frequentist binomial GLMM with subpopulation as random factor and the brown bear *Occurrence Database* as presences/pseudo-absences (Supplementary Table 3, Supplementary Figure 1). We compared the univariate models using Akaike Information Criterion (AIC)^79^.

### Brown bear presence databases

We constructed two databases of brown bear presence: (1) a database of brown bear at the scale of its geographic range using historical distribution data, the *Range Database*, and (2) a database of brown bear using current data, the *Occurrence Database*.

The *Range Database* contains data, at a low spatial resolution (10 x 10 km), on the Eurasian historical distribution of brown bear, areas of current presence and areas which were occupied in the past but where the species has been extirpated. We discarded the North American brown bear distribution due to large differences in life-history traits between Eurasian and North American subspecies^27, 88^. We used the IUCN historical distribution of the brown bear^28^ and added information on past distributions from several sources of published data and historical references^89–94^ (Supplementary Figure 13). The brown bear is a species that has suffered intense human persecution resulting in multiple local extinctions over the last two millennia^95^. Thus, we included information of past brown bear distribution since the 1st century, even though human-induced local extirpation of brown bears could have occurred before, e.g., in areas surrounding the Mediterranean region^96^. Absences were extracted randomly within a 50 km buffer around pixels with brown bear presences in the historical distribution, and we selected the same number of pseudo-absences as presences. Across the pixels of presences and absences we applied a stratified sampling procedure following section 7.4.3 in Guisan et al. 2017^97^: first we used climate variables (*Clim_3*, *Clim_4*, *Clim_8* and *Clim_10*) to create stratums and then we selected an equal number of presences and absences by stratum. This last selection was used to model the distribution of brown bear at the range scale.

The *Occurrence Database* contains brown bear occurrences in Europe and Turkey with an uncertainty of < 1 km^2^ for the period 1989–2018. We used data from (a) systematic surveys, e.g., excluding locations of occasional sightings of bear damage, hunted bears and road/train casualties, and (b) bears equipped with GPS/VHF collars, from which we excluded bears that were intentionally trapped and collared to be permanently monitored due to their problematic behaviour (e.g., bears selected because they frequently approach human settlements, or frequently attack livestock and/or cause other types of damage) to avoid bias in the data towards problematic bears, but we did not exclude randomly trapped bears that exhibited problematic behaviour. The resulting database contained data from 37 different research groups, comprising more than 70 researchers, and included data from all European and Turkish subpopulations and from all countries, except Belarus, with a stable presence of brown bears. The database was formed from 52 original datasets with more than 3.2 million occurrences from diverse sources: 97.56% from GPS collars, 0.53% from VHF collars, 1.12% from tracks, 0.01% from camera traps and 0.01% from unspecified sources (Supplementary Tables 1 and 2). Occurrences were rasterized to a binary map of brown bear occurrence (presences = 1) and non-occurrence (pseudo-absences = 0). We obtained more than 100,000 pixels with the presence of brown bear showing a spatial bias between the different subpopulations, with a minimum of 471 presences for the Turkey subpopulation and a maximum of 45,119 presences for the Scandinavian subpopulation (Supplementary Table 3 and Supplementary Figure 1). For models using the *Occurrence Database*, we first excluded data occurring in non-terrestrial systems (i.e., presences in lakes or seas). Then, to avoid overrepresentation of some subpopulations due to a larger number of individuals or greater sampling effort, and also because of computing limitations, we selected a maximum of 2,000 presences for each subpopulation. For subpopulations with less than 2,000 presences available, we selected all presences available (Supplementary Table 3). Pseudo-absences were extracted randomly within a 5 km buffer around pixels with brown bear presences, and we selected the same number of pseudo-absences as presences for each subpopulation. In total, we used 24,908 presences, which were split into two sets, one set to train the model (19,926 presences, 80%) and the other to validate the model (4,982 presences, 20%). The training data were used in all models using the *Occurrence Database* (for comparison of biotic proxies explaining brown bear distribution and for modelling brown bear distribution at a fine scale with the Bayesian models, BMs).

### Bayesian brown bear species distribution model

#### Species distribution model of the historical range

Data at the range scale are of low resolution but larger extent, which capture a greater variability of the effect on species presence of factors acting at larger scales, e.g., climate, and avoid bias or the truncation of the environmental space^33, 34^. Data at the habitat scale are of high resolution but small extent, which improve spatial precision and our understanding of the effect on species presence of factors acting at smaller scales. As we know that the brown bear distribution has contracted as a result of human activity, we wanted to explore how the historical relationship of brown bear occurrences with climate variables might influence our distribution models. For example, if the brown bear historically thrived in warmer conditions, then this information could also help us better predict its current and future distribution by avoiding a truncation of environmental variables^34^. To this end, we used a model of the historical distribution of the brown bear (based on historical range data and climate data) to inform models of the current distribution. Specifically, we fitted a species distribution model with presences/absences from the *Range Database* (10 x 10 km resolution) as a function of historical bioclimatic variables (Supplementary Appendix 2). We selected bioclimatic variables by first dropping those with a VIF over a threshold of 10 using the R package usdm^58, 97^. We further refined the variables by selecting the four best historical climate variables (Supplementary Table 61) on the basis of the AIC of univariate binomial GLMs (logit link) with linear and quadratic effects. We then fitted a final ‘historical’ distribution model with a Bayesian binomial GLM using these four variables (*Clim_3*, *Clim_4*, *Clim_8* and *Clim_10*) and a stratified sampling of presences/absences from the brown bear *Range Database*. We extracted the mean and standard deviation of the parameter estimates for use as an informed prior for the models described below (Supplementary Tables 62-66 and Supplementary Figures 14-17).

#### Modelling and predicting brown bear distribution at a fine scale

We modelled the distribution of the brown bear at a high spatial resolution (1*×*1 km; Supplementary Table 42) using as response variable the training selection of presences/pseudo-absences from the brown bear *Occurrence Database* (N presences = 19,926 ; N pseudo-absences = 19,926; Supplementary Tables 1-3) and three types of predictor variables: climate, land-use and biotic. We selected four variables to include within each type. For climate variables, we selected the same four variables used in the model of the historical brown bear distribution (but with values for the current climate, obtained from the CHELSA database^29^). For land-use and biotic variables, we selected the four best (AIC-based) uncorrelated variables determined by a univariable GLMM (Supplementary Tables 67, 68 and 40).

We fitted three Bayesian models (BMs) explaining brown bear distribution using different combinations of factors (abiotic, biotic or both): (1) a model with abiotic (climate and land use variables) and biotic predictors (using biotic variables), (2) a model with abiotic predictors only, and (3) a model with biotic predictors only (Supplementary Table 41). Note that only models including climate variables can use the historical priors, and thus the model based solely on biotic variables can only use current data. We also fitted a null model with only the intercept using only current presences/pseudo-absences.

The two Bayesian hierarchical models (BHMs) which used historical range data (the model with abiotic and biotic predictors and the model with biotic predictors only), used non-informative priors for land use and biotic variables, and informative priors based on the historical distribution model as defined above. We assumed equal confidence for the historical and current data, and thus the mean and SD of the parameter estimates from the historical distribution model were not modified and used directly as priors. The biotic model used non-informative priors for all variables. All models were Bayesian binomial GLMMs calculated with a logit link, using the Hamiltonian Monte Carlo algorithm in Stan (mc-stan.org). We used Stan with the rstan, rtanarm and loo packages in R to fit and assess the diagnostics of the models^98–100^. We evaluated and compared the models using the Widely Applicable Information Criterion (WAIC)^101^. The best model (based on the WAIC) was evaluated using the validation subset of the brown bear *Occurrence Database* (N presences = 4,982; N pseudo-absences = 4,982; Supplementary Table 41). As we used pseudo-absences, we established a cut-off for the potential distribution based on the 90th percentile training presence^102, 103^, that is, leaving out 10% of the observed presences of the training dataset.

In addition to those models, and in order to evaluate whether there was an effect of combining different data, we fitted two simple Bayesian models (not hierarchical) for the two BHMs without considering the information of the historical range, we used non-informative priors for all predictors (Supplementary Table 42).

We used the best model to predict the distribution of brown bear in all subpopulations for the current and nine future climate/land use change scenarios which considered combinations of three SSPs (SSP1-2.6, SSP3-6.0 and SSP5-8.5) with changes in (1) abiotic and biotic variables, (2) abiotic variables and (3) biotic variables. For the biotic variables, the climate and land use scenarios were used indirectly to predict the influence of climate and land use on the habitat of species in the brown bear diet, which was then summarised as energy available in the space as described above. For the current and each of the nine future climate/land use change scenarios for the distribution of brown bear, we calculated different descriptors related to the conservation status of species^104, 105^, such as area of the distribution, percentage of the distribution occupied,and distribution included in protected areas using the World Database of Protected Areas^106^. All statistical analyses were performed in the R program^107^.

## ACKNOWLEDGEMENTS

PML, LJP, WT, NS, MDB, LM, NBa, TD, AF, SCF, AZ were financially supported through the 2015-2016 BiodivERsA COFUND call, with national funders Agence Nationale de la Recherche (ANR), France (Grant number: ANR-16-EBI3-0003), National Science Center (NCN), Poland (Grant number: 2016/22/Z/NZ8/00121), Federal Ministry of Education and Research (BMBF) and DLR Project Management Agency (DLR-PT), Germany (Grant number: 01LC1614A), Romanian National Authority for Scientific Research and Innovation, CCCDI – UEFISCDI, Romania (Grant number: BiodivERsA3-2015-147-BearConnect (96/2016) within PNCDI III, and Norwegian Research Council (RCN), Norway (Grant number: 269863). VPe was financially supported by the Grant PID2020-114181GB-I00 funded by MCIN/AEI/ 10.13039/501100011033 and by “ERDF A way of making Europe”. DZ and AD were funded by the PIN-MATRA and the BBI-MATRA programs and the project LIFE07NAT/IT/000502. NBo and DC were funded by the Ministry of Education Science and Technological Development of the Republic of Serbia (451-03-68/2022-14-200007) and by the WWF Adria. SF and LF were funded by the Regione Friuli Venezia Giulia. DH, SR and DDA thanks the funding from LIFE DINALP BEAR and EURONATUR. KJ and TS acknowledge the funding from the LIFE DINALP BEAR (LIFE13 NAT/SI/000550) and the Slovenia Research Agency, project J4-7362. AM-J and PMo were funded by WWF Austria, LIFE13 NAT/SI/000550 LIFE DINALP BEAR. AAK and MdGH thanks the funding to Vodafone, Acturos and the Hellenic Ministry of Rural Development. JK and HB thanks the funding from the Swedish Environmental Protection Agency, Norwegian Environmental Agency. YM and MPs were funded by the LIFE program (LIFE07 NAT/GR/000291; LIFE07 NAT/IT/000502; LIFE12 NAT/GR/000784; LIFE09 NAT/GR/000333; LIFE15 NAT/GR/001108; LIFE96NAT/GR/003222; LIFE99NAT/GR/006498). ER and JN thanks the funding from the Principado de Asturias government. I-MP was funded by LIFE08NAT/RO/000500- LIFEURSUS (financed by LIFE+ and ACDB) and by ROSCI0229 Siriu (Romanian Environmental SOP 2007-2014). US and ET thanks the funding to the Institutional research funding (IUT20-32) and the grant PRG1209 from the Estonian Ministry of Education and Research. NS, TZ-K, ASe, FZ thanks the funding from Project GLOBE (POLNOR/198352/85/2013) funded by the Polish-Norwegian Research Programme operated by the National Centre for Research and Development and by the budget of Tatra National Park. ASo, AE thanks the fuding from Hacettepe University, Kastamonu University, General Directorate of Nature Conservation and National Parks, Ministry of Forestry and Water Affairs, Turkey. ASt and DM were funded by the the Balkan Lynx Recovery Programme. SCF and AZ thanks the funding from The Scandinavian Brown Bear Research Project, which is funded by the Swedish Environmental Protection Agency, the Norwegian Directorate for Nature Management and the Austrian Science Fund. CCB and HA were funded by Kaçkar Mountains Sustainable Forest Use and Conservation Project. IK, SH and OH were funded by Ministry of Agriculture and Forestry, Government of Finland. MPo, RJ, GI, OI, and GS were funded by Life for Bear LIFE 13 NAT/RO/001154. the Nulceu Programmme ANCSI Eco-etologa carnicorelor mari in contextual dezvoltarii infrastructurii, and the BEAR Ethology Around Romania FP5. AOS and DO were funded by the Ministry of Forestry and Water Affairs from Turkey, the Nature Conservation Centre and the United Nations GEF-5 Programme, by the Scientific and Technological Research Council of Turkey. CHS and MWC were funded by the Christensen Fund, National Geographic Society, UNDP Small Grants Programme, University of Utah, Whitley Fund. APS, VNP were funded by the Russian Science Foundation (project No. 18-14-00093). AT and BH were funded by MAVA foundation, EuroNatur foundation. CD was funded by Milvus Group’s, Bears in Mind (the Netherlands), Bernd Thies Foundation (Switzerland), Columbus Zoo and Aquarium (USA), EuroNatur (Germany), Frankfurt Zoological Society (Germany), the International Association for Bear Research and Management (USA) and the Nando Peretti Foundation (Italy). We thank Manuela Gonzalez-Suarez, Pablo Burraco and Sören Faurby for their helpful suggestions regarding the manuscript. We are also in debt to Angel Lucas for the artwork in Figs. 1, 3–5. We also thank Galician Supercomputing Center (CESGA, Galicia, Spain) for their technical and infrastructure support. We thank GBIF and people and institutions that contribute to GBIF, to make public their environmental research data. We would like to thank to the Brown Bear Foundation rangers and technicians, regional governments of Cantabria, Galicia, Asturias, Cantabria and Castilla y León, Regione Friuli Venezia Giulia, the ISPRA, the Slovenia Forest Service, the Hunters Society of Slovenia, the CUFAA, the University of Thessaloniki, the Northern Pindos National Park, the AnGre Regional Development, the AnKas Regional Development, the Rodopi Mountain Range National Park, the University of Thessaly, Egnatia SA, Ministry of Environment, Energy and Climatic Change, Government of Navarra, Government of Aragon, Government of Andorra, Conselh Generau d’Aran, Fauna and Flora Service, Alt Pirineu Natural Park, Ramón Jato, Ivan Afonso, Sergio Mir, Salvador Gonçalbez, Antoni Batet, Jordi Guillén, Xavier Garreta, members and students of the Carpathian Brown Bear Project, the personnel of Tatra National Park helping in the captures, the Hacettepe University, the Regional Inspectorate of Environment and Waters – Smolyan, the Central Balkan National Park, the Vitosha Nature park, Sanna Kokko, the Natural Resources Institute Finland, to Slavomír Finďo, the WWF Austria, the Institute of Wildlife Biology and Game Management, the University of Natural Resources and Life Sciences, Vienna, to Eko-Zon Public Health and Environmental Consultation, to Kolesnikov V.V., Mashkin V.I., Skumatov D.V., Zarubin B.E. (all – VNIIOZ, Kirov), the Balkan Lynx Recovery Programme, the DVM Levente Borka-Vitális (Vets4Wild), Károly Illyés (“MureLul” Private Forestry Service), Károly Pál (“Pro Diana” Hunting Association) and Ágoston Pál (“Târnava Mare” Hunting and Angling Association).

## AUTHOR CONTRIBUTIONS

PML led the conceptualization under the advice of WT and LJP; PML led the data curation, including the construction of the brown bear *Occurrence Database*; PML led the construction of the *Trophic Database* with contributions from JA and NS; NS suggested to reassign rEDEC and modified it with JA and PML; PML led the classification of wild/food species with contributions from NS, MDB, LM, HB, JK and SO; DDA led the work in the *Range Database* with contributions from JN and PML; PML led the statistical analysis with input from WT, MVT, EP, JA, MG, ER and LJP; WT, JA, NS, MDB, LM, NBa, AF and AZ acquired the funding; PML, JA, NS, LM, SCF, AZ, IA-J, HA, FB, A-TB, CCB, NBo, EB, KB, NBr, HB, MWC, DC, PC, AC, DDA, MdGH, CD, AD, AE, SF, LF, CG, SH, BH, DH, OH, GI, OI, KJ, IK, JVL-B, PMä, DM, YM, PMo, AM-J, AM, JN, SO, DO, SP, LPe, AP, VNP, I-MP, MPo, MPs, P-YQ, GR, SR, ER, US, APS, AOS, CHS, ASe, GS, TS, MS, ASo, ASt, ET, KT, AT, IT, TT, FZ, DZ, TZ-K collected data; PML led the methodology with input from WT, MVT, EP, JA, LM, MG, and LJP; NS and MDB contributed with project administration tasks; WT, MVT, MG and LJP provided resources; PML led the programming/coding with input from WT, MVT, EP, JA, and MG; WT and LJP supervised the results and research outputs; PML led the preparation of figures with input from WT, MVT, EP, JA, VPe and LJP; PML wrote the original draft with input from WT, MVT, EP, VPe and LJP; PML led the review and preparation of the manuscript with input from WT, MVT, EP, JA, NS, MDB, LM, VPe, MG, TD, AF, SCF, HA, KB, AC, DDA, CD, SF, LF, DH, OH, KJ, JK, IK, JVL-B, AM-J, ASo, ASt, TS, ASe, AT, DZ, LJP. PML coordinated with all coauthors/stakeholders to get brown bear occurrences. All authors read and approved the final version of the manuscript.

## SUPPLEMENTARY INFORMATION

The online version contains supplementary material.

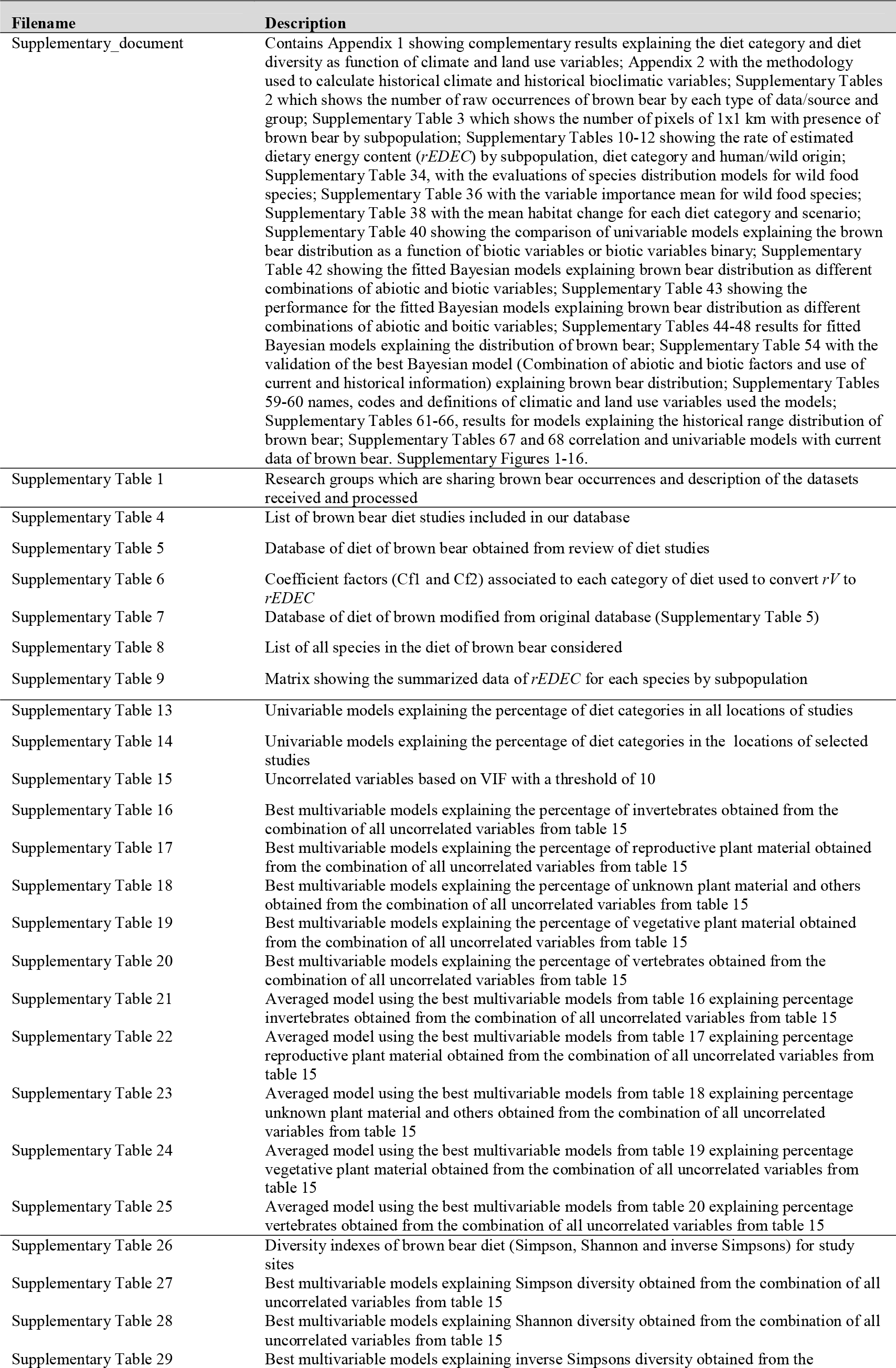
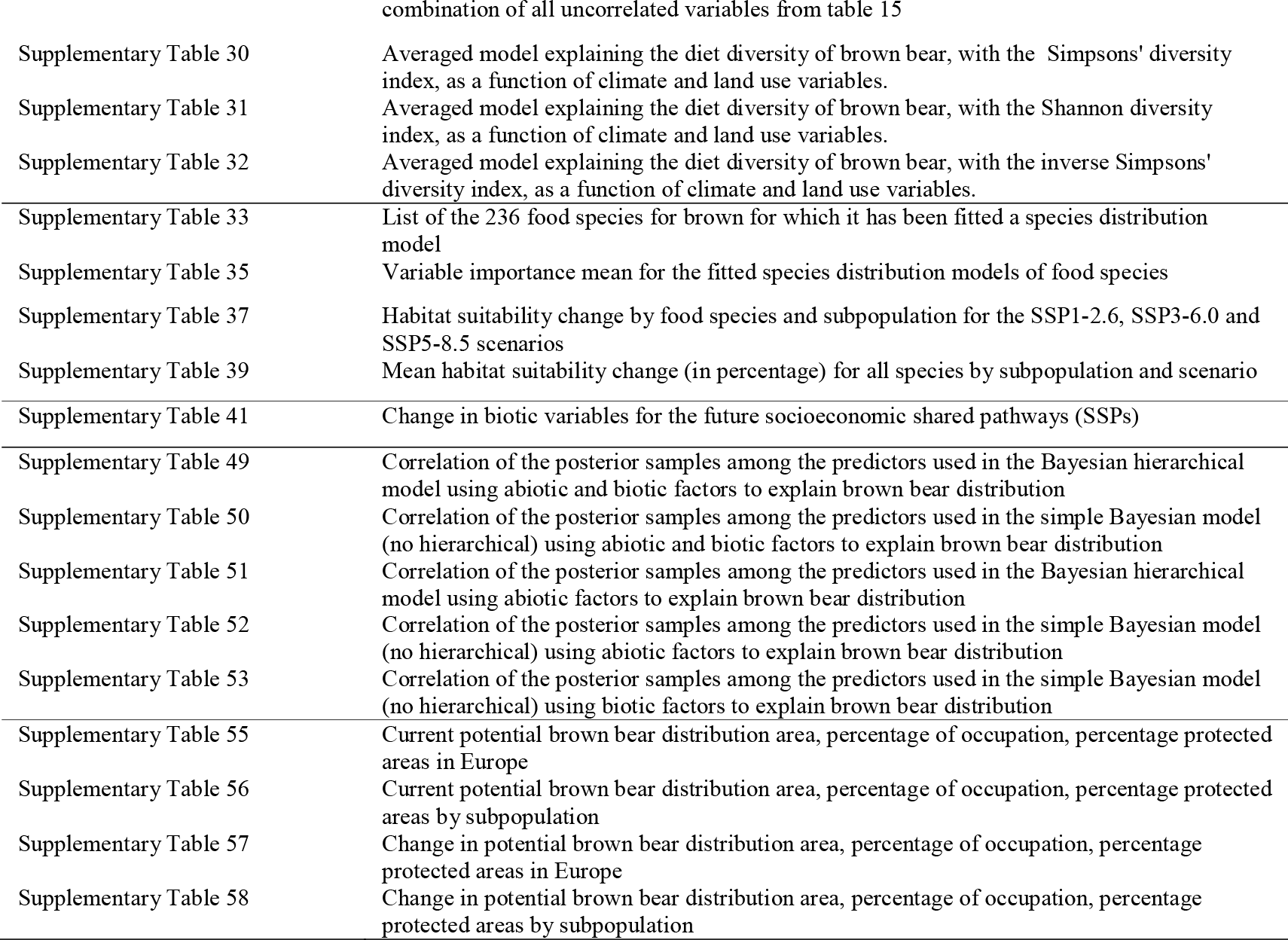

