## Supplementary_document for "Including biotic interactions in species distribution models improves the understanding of species niche: a case of study with the brown bear in Europe"

PM Lucas: Institute of Nature Conservation, Polish Academy of Sciences, Adama Mickiewicza 33, 31-120 Kraków, Poland. Department of Biology and Biotechnologies “Charles Darwin”,

Sapienza Università di Roma, Viale dell'Università 32, 00185 Rome, Italy. ORCID: 0000-0003-4517-9748.

W Thuiller: Univ. Grenoble Alpes, Univ. Savoie Mont Blanc, CNRS, LECA, F-38000 Grenoble, France. ORCID: 0000-0002-5388-5274.

MV Talluto: Department of Ecology, University of Innsbruck, Technikerstraße 25, 6020 Innsbruck, Austria. ORCID: 0000-0001-5188-7332.

E Polaina: Department of Ecology, Swedish University of Agricultural Sciences, 75007 Uppsala, Sweden. ORCID: 0000-0002-5064-5881.

J Albrecht: Senckenberg Biodiversity and Climate Research Centre (SBiK-F), Senckenberganlage 25, 60325 Frankfurt am Main, Germany. ORCID: 0000-0002-9708-9413.

N Selva: Institute of Nature Conservation, Polish Academy of Sciences, Adama Mickiewicza 33, 31-120 Kraków, Poland. Departamento de Ciencias Integradas, Facultad de Ciencias Experimentales, Centro de Estudios Avanzados en Física, Matemáticas y Computación, Universidad de Huelva, 21071 Huelva, Spain. ORCID: 0000-0003-3389-201X.

M De Barba: Univ. Grenoble Alpes, Univ. Savoie Mont Blanc, CNRS, LECA, F-38000 Grenoble, France. Biotechnical Faculty, Department of Biology, University of Ljubljana, Jamnikarjeva 101, SI-1000, Ljubljana, Slovenia. DivjaLabs Ltd., Aljaževa ulica 35a, 1000 Ljubljana, Slovenia. ORCID: 0000-0002-2979-3716.

L Maiorano: Department of Biology and Biotechnologies "Charles Darwin", Sapienza Università di Roma, Viale dell'Università 32, 00185 Rome, Italy. ORCID: 0000-0002-2957-8979.

V Penteriani: Department of Evolutionary Ecology, National Museum of Natural Sciences (MNCN-CSIC), Madrid, Spain. ORCID: 0000-0002-9333-7846.

M Guéguen: Univ. Grenoble Alpes, Univ. Savoie Mont Blanc, CNRS, LECA, F-38000 Grenoble, France. ORCID: 0000-0002-1045-2997.

N Balkenhol: Wildlife Sciences, University of Goettingen, Buesgenweg 3, 37077 Goettingen, Germany. ORCID: 0000-0003-4921-5443.

T Dutta: Wildlife Sciences, University of Goettingen, Buesgenweg 3, 37077 Goettingen, Germany. European Forest Institute, Platz der Vereinten Nationen 7, 53113 Bonn, Germany. ORCID: 0000-0002-5236-2658.

A Fedorca: Department of Wildlife, National Institute for Research and Development in Forestry “Marin Drăcea”, Closca 13, Brasov, Romania. Department of Silviculture, Transilvania University of Brasov, Beethoven Line 1, Brasov, Romania. ORCID: 0000-0001-5828-5422.

SC Frank: Faculty of Technology, Natural Sciences and Maritime Sciences, Department of Natural Sciences and Environmental Health, University of South-Eastern Norway, 3800 Bø i Telemark, Norway. ORCID: 0000-0001-8153-6656.

A Zedrosser: Faculty of Technology, Natural Sciences and Maritime Sciences, Department of Natural Sciences and Environmental Health, University of South-Eastern Norway, 3800 Bø i Telemark, Norway. ORCID: 0000-0003-4417-3037.

I Afonso-Jordana: Departament de Territòri, Paisatge e Gestion Ambientau, Conselh Generau d'Aran, Plaça d'Aran 1-2, 25530 Vielha, Spain.

H Ambarlı: Department of Wildlife Ecology and Management, Faculty of Forestry No: 130, Düzce University, 81620, Düzce, Turkey. ORCID: 0000-0003-4336-9417.

F Ballesteros: Brown Bear Foundation, C/ San Luis 17, 39010, Santander, Spain. ORCID: 0000-0003-4764-4041.

A-T Bashta: Institute of Ecology of the Carpathians NAS Ukraine, Kozelnytska st. 4, 79026 Lviv, Ukraine. ORCID: 0000-0002-8134-5507.

CC Bilgin: Department of Biology, Middle East Technical University, 06800 Ankara, Turkey. ORCID: 0000-0001-9284-307X.

N Bogdanović: Faculty of Biology, University of Belgrade, Studentski trg 16, 11000 Belgrade, Serbia. ORCID: 0000-0002-3782-6602.

E Bojārs: Estonian University of Life Sciences, Fr.R.Kreutzwaldi 1, 51006 Tartu, Estonia.

K Bojarska: Institute of Nature Conservation, Polish Academy of Sciences, Adama Mickiewicza 33, 31-120 Kraków, Poland. ORCID: 0000-0001-7872-5763.

N Bragalanti: Servizio Foreste e Fauna, Provincia Autonoma di Trento, Via Giovanni Battista Trener 3, 38121 Trento, TN, Italy.

H Brøseth: Norwegian Institute for Nature Research (NINA), NO-7485 Trondheim, Norway. ORCID: 0000-0003-3795-891X.

MW Chynoweth: Department of Wildland Resources, Utah State University, Uintah Basin, 320 North Aggie Blvd., Vernal, UT 84078, USA. ORCID: 0000-0001-9203-8193.

D Ćirović: Faculty of Biology, University of Belgrade, Studentski trg 16, 11000 Belgrade, Serbia. ORCID: 0000-0001-9468-0948.

P Ciucci: Department of Biology and Biotechnologies "Charles Darwin", Sapienza Università di Roma, Viale dell'Università 32, 00185 Rome, Italy. ORCID: 0000-0002-0994-3422.

A Corradini: Department of Civil, Environmental and Mechanical Engineering (DICAM), University of Trento, via Mesiano 77, 38123 Trento, TN, Italy. Animal Ecology Unit, Research and Innovation Centre, Fondazione Edmund Mach, via Mach 1, 38010 San Michele all'Adige, TN, Italy; Stelvio National Park, Via De Simoni 42, 23032 Bormio, SO, Italy. ORCID: 0000-0002-9294-656X.

D De Angelis: Department of Biology and Biotechnologies "Charles Darwin", Sapienza Università di Roma, Viale dell'Università 32, 00185 Rome, Italy. ORCID: 0000-0002-2529-6210.

M de Gabriel Hernando: Department of Conservation Biology, Estación Biológica de Doñana (EBD-CSIC), Avd. Américo Vespucio 26, 41092, Sevilla, Spain. ORCID: 0000-0002-8722-6146.

C Domokos: Milvus Group Bird and Nature Protection Association, Crinului 22, 540343 Tîrgu Mureș, Romania. ORCID: 0000-0002-9973-7779.

A Dutsov: Balkani Wildlife Society, 67 Tsanko Tserkovski str, Entr.3, floor 2, apt.3, BG-1421 Sofia, Bulgaria.

A Ertürk: Hunting and Wildlife Program, Kastamonu University, 37800, Araç, Kastamonu, Turkey. ORCID: 0000-0001-5498-3856.

S. Filacorda: Department of Agri-Food, Environmental and Animal Sciences, University of Udine, Via Sondrio 2/A, 33100 Udine, Italy. ORCID: 0000-0002-0984-2373.

L Frangini: Department of Agri-Food, Environmental and Animal Sciences, University of Udine, Via Sondrio 2/A, 33100 Udine, Italy. ORCID: 0000-0002-1281-0188.

C Groff: Servizio Foreste e Fauna, Provincia Autonoma di Trento, Via Giovanni Battista Trener 3, 38121 Trento, TN, Italy.

S Heikkinen: Natural Resources Institute Finland, Latokartanonkaari 9, 00790 Helsinki, Finland.

B Hoxha: Protection and Preservation of Natural Environment in Albania, Rr. Janos Hunyadi P.32/11, 1019 Tirana, Albania.

D Huber: Faculty of Veterinary Medicine, University of Zagreb, Heinzelova 55, 10000 Zagreb, Croatia. ORCID: 0000-0002-3912-3812.

O Huitu: Natural Resources Institute Finland, Latokartanonkaari 9, 00790 Helsinki, Finland. ORCID: 0000-0003-3829-1213.

G Ionescu: Department of Wildlife, National Institute for Research and Development in Forestry "Marin Drăcea, " Closca 13, Brasov, Romania.

O Ionescu: Department of Wildlife, National Institute for Research and Development in Forestry "Marin Drăcea, " Closca 13, Brasov, Romania.

K Jerina: Department for Forestry, Biotechnical Faculty, University of Ljubljana, Večna pot 83, 1000 Ljubljana, Slovenia. ORCID: 0000-0003-3946-2678.

R Jurj: Department of Wildlife, National Institute for Research and Development in Forestry "Marin Drăcea, " Closca 13, Brasov, Romania.

AA Karamanlidis: ARCTUROS, Civil Society for the Protection and Management of Wildlife and the Natural Environment, Florina, Greece; Faculty of Environmental Sciences and Natural Resource Management, Norwegian University of Life Sciences, Ås, Norway. ORCID: 0000-0003-0943-1619.

J Kindberg: Norwegian Institute for Nature Research, Trondheim, Norway and department of Wildlife, Fish and Environmental Studies, Faculty of Forest Sciences, Swedish University of Agricultural Sciences, Umeå, Sweden. ORCID: 0000-0003-1445-4524.

I Kojola: Natural Resources Institute Finland, Latokartanonkaari 9, 00790 Helsinki, Finland. ORCID: 0000-0003-2866-5090.

JV López-Bao: Biodiversity Research Institute (CSIC - Oviedo University - Principality of Asturias), Oviedo University, 33600 Mieres, Spain. ORCID: 0000-0001-9213-998X.

P Männil: Estonian Environment Agency, Mustamäe tee 33, 10616 Tallinn, Estonia.

D Melovski: Macedonian Ecological Society, Arhimedova 5, 1000 Skopje, N.Macedonia; Wildlife Sciences, University of Goettingen, Busgenweg 3, 37077 Goettingen, Germany. ORCID: 0000-0002-1072-6978.

Y Mertzanis: "Callisto" Wildlife and Nature Conservation Society, Greece and Department of Ecology, School of Biology, Aristotle University, Thessaloniki, Greece.

P Molinari: Progetto Lince Italia, Via Roma 43, I-33018 Tarvisio, Italy.

A Molinari-Jobin: Progetto Lince Italia, Via Roma 43, I-33018 Tarvisio, Italy. ORCID: 0000-0002-3729-2532.

A Mustoni: Adamello-Brenta Natural Park, Via Nazionale 12, 38080 Strembo, Italy.

J Naves: Department of Conservation Biology, Estación Biológica de Doñana (EBD-CSIC), Avd. Américo Vespucio 26, Sevilla, 41092, Spain. ORCID: 0000-0003-3773-0288.

S Ogurtsov: Central Forest State Nature Biosphere Reserve, Zapovedniy, Tver region, 172521 Russian Federation. ORCID: 0000-0002-0859-8954.

D Özüt: Nature Conservation Centre, Aşağı Öveçler Mah. 1065. Cad. 1293 Sok. No. 9/32, 06460 Ankara, Turkey. ORCID: 0000-0002-6692-7533.

S Palazón: Fauna and Flora Service, Generalitat of Catalonia. Provença 204-208, 08036 Barcelona, Spain.

L Pedrotti: Stelvio National Park, Via De Simoni 42, 23032 Bormio, SO, Italy; Servizio Faunistico, Provincia Autonoma di Trento, Via Trener 3, 38121 Trento, TN, Italy.

A Perović: Centre for protection and research of birds of Montenegro - CZIP, Velje brdo 35, 81412 Podgorica, Montenegro.

V Piminov: Department of Game Resources, Russian Research Institute of Game Management and Fur Farming, 79 Preobrazhenskaya Str., Kirov, 610000, Russian Federation.

I-M Pop: Centre for Environmental Research and Impact Studies (CCMESI), University of Bucharest, 1 N. Balcescu Blvd., Bucharest, Romania. The Association for the Conservation of

Biological Diversity (ACDB), 12 Ion Creanga St., Focsani, Romania. ORCID: 0000-0001-9706-3697.

M Popa: Department of Wildlife, National Institute for Research and Development in Forestry “Marin Drăcea”, Closca 13, Brasov, Romania. Department of Silviculture, Transilvania University of Brasov, Beethoven Line 1, Brasov, Romania.

M Psaralexi: “Callisto” Wildlife and Nature Conservation Society, Greece and Department of Ecology, School of Biology, Aristotle University, Thessaloniki, Greece. ORCID: 0000-0002-0220-8855.

P-Y Quenette: Research and Scientific Support Direction, French Biodiversity Agency, Impasse de la Chapelle, 31800 Villeneuve de Rivière, France.

G Rauer: WWF Austria, Ottakringer Strasse 114-116, 1160 Vienna, Austria. Institute of Wildlife Biology and Game Management, University of Natural Resources and Life Sciences, Gregor-Mendel-Haus Gregor-Mendel-Straße 33, 1180 Vienna, Austria.

S Reljic: Faculty of Veterinary Medicine, University of Zagreb, Heinzelova 55, 10000 Zagreb, Croatia. ORCID: 0000-0002-8110-384X.

E Revilla: Department of Conservation Biology, Estación Biológica de Doñana (EBD-CSIC), Avd. Américo Vespucio 26, Sevilla, 41092, Spain. ORCID: 0000-0001-5534-5581.

U Saarma: Department of Zoology, Institute of Ecology and Earth Sciences, University of Tartu, J. Liivi 2, 50409 Tartu, Estonia. ORCID: 0000-0002-3222-571X.

AP Saveljev: Department of Animal Ecology, Russian Research Institute of Game Management and Fur Farming, 79 Preobrazhenskaya Str., Kirov, 610000, Russian Federation. ORCID: 0000-0002-8103-5787.

AO Sayar: Department of Game and Wildlife, Cankiri Karatekin University, Fatih, Uluyazı Kampüsü Ring Yolu, 18100 Çankırı Merkez/Çankırı, Turkey. ORCID: 0000-0001-6000-2946.

CH Şekercioğlu: School of Biological Sciences, University of Utah, Salt Lake City, UT, United States. Department of Molecular Biology and Genetics, Koç University, Sarıyer, İstanbul, Turkey. ORCID: 0000-0003-3193-0377.

A Sergiel: Institute of Nature Conservation, Polish Academy of Sciences, Adama Mickiewicza 33, 31120 Krakow, Poland. ORCID: 0000-0002-3455-4218.

G Sîrbu: Department of Wildlife, National Institute for Research and Development in Forestry “Marin Drăcea”, Closca 13, Brasov, Romania. Department of Silviculture, Transilvania University of Brasov, Beethoven Line 1, Brasov, Romania.

T Skrbinšek: Biotechnical Faculty, Department of Biology, University of Ljubljana, Jamnikarjeva 101, SI-1000, Ljubljana, Slovenia. DivjaLabs Ltd., Aljaževa ulica 35a, 1000 Ljubljana, Slovenia. ORCID: 0000-0003-4435-7477.

M Skuban: Carpathian Widlife Society, Tulská 2461/29, 96001 Zvolen, Slovakia.

A Soyumert: Hunting and Wildlife Program, Kastamonu University, 37800, Araç, Kastamonu, Turkey. ORCID: 0000-0003-0196-9617.

Aleksandar Stojanov: Macedonian Ecological Society, Arhimedova 5, 1000 Skopje, North Macedonia.

E Tammeleht: Department of Zoology, Institute of Ecology and Earth Sciences, University of Tartu, J. Liivi 2, 50409 Tartu, Estonia. ORCID: 0000-0003-0871-051X.

K Tirronen: Institute of Biology, Karelian Research Centre, Russian Academy of Sciences, 11 Pushkinskaya Str. Petrozavodsk, 185910 Russian Federation. ORCID: 0000-0002-8208-5460.

A Trajçe: Protection and Preservation of Natural Environment in Albania, Rr. Janos Hunyadi P.32/11, 1019 Tirana, Albania. ORCID: 0000-0002-6815-7421.

I Trbojević: Faculty of Natural Science and Mathematics, University of Banja Luka, Dr. Mladena Stojanovića 2, 78000 Banja Luka, Bosnia and Herzegovina; Faculty of Ecology, Independent University of Banja Luka, Veljka Mladjenovića 12e, 78000 Banja Luka, Bosnia and Herzegovina. ORCID: 0000-0003-0781-7235.

T Trbojević: Ecology and Research Association, Dr. Mladena Stojanovića 2, 78000 Banja Luka, Bosnia and Herzegovina.

F Zięba: Tatra National Park, Kuźnice 1, 34-500 Zakopane, Poland.

D Zlatanova: Faculty of Zoology, Sofia University “St. Kliment Ohridski”, Bul. Dragan Tsankov 8, 1164 Sofia, Bulgaria. ORCID: 0000-0002-4115-2786.

T Zwijacz-Kozica: Tatra National Park, Kuźnice 1, 34-500 Zakopane, Poland. ORCID: 0000-0002-7488-975X .

LJ Pollock: Department of Biology, McGill University, 1205 Docteur Penfield, Montréal, Canada H3A 1B1.

### Appendix 1

**Associations between diet and environmental variables**

Among climate variables, the mean diurnal temperature range (*Clim_2*) had the most important role in explaining the share of different food categories in brown bear dietary energy content (Supplementary Tables 21-25). In areas with higher diurnal temperature range, bears consumed less vegetative plant material, less unknown plant material and fewer vertebrates. The precipitation affected bear feeding habits to a lesser extent than temperature. The share of the food category “unknown plant material and others” in bear dietary content was positively related to the mean temperature of the warmest quarter, the seasonality of the precipitation (*Clim_15*) and to mean diurnal temperature range, while it correlated negatively with the precipitation of the wettest month and to annual temperature range (*Clim_7*; Supplementary Tables 13-32).

Among land use variables, the percentage of broadleaf forests (*LC_3*), of bare areas (*LC_9*) and of sparse vegetation (*LC_7*) were the most important in shaping the brown bear dietary energy content. In turn, in areas where bare areas were common, bears ingested more invertebrates but less reproductive plant parts. The proportion of rangeland (*LC_8*) had the most significant impact on the brown bear dietary content; it related positively to the share of vertebrates and vegetative plant material, as well as the overall dietary diversity. Cultivated areas and pastures (*LC_2*) negatively affected the consumption of vertebrates and vegetative plant material. The urban areas (*LC_1*) affected only, and tentatively, the percentage of vegetative plant parts in the bear diet. The proportion of the food category “unknown plant material and others” in bear diet related positively to the percentage of natural landscape (*Nat. Landscape*), urban areas (*LC_1*) and bare habitats (*LC_9*), and negatively to broadleaf forest (*LC_3*) and rangeland (*LC_8*;

Moreover, bears had a more diverse dietary content in areas with lower mean diurnal temperature range (*Clim_2*), and a less cover of broadleaved forest (*LC_3*). The diet diversity was, however, positively related to the percentage of rangeland areas (*LC_8*). In areas where broadleaf forests had higher contribution to land cover, bears displayed a more diverse diet (Supplementary Tables 13-32).

### Appendix 2

**Calculation of historical bioclimatic variables**

We used the Time-Series (TS) Version 3.21 provided by the Climate Research Unit[108] at a resolution of 30 arcminutes. The data provide monthly values for different climatic conditions over the last century. The IPCC defines climate as the mean and variability of weather variables over a period of time, which can range from months to millions of years[109]. We used the classical range of 30 years adopted by the World Meteorological Organization[110] and selected as historical climate the period from 1901 to 1931 to calculate representative monthly precipitation, minimum temperature and maximum temperature of that period. Using package dismo[111] in R we calculated the bioclimatic variables.

### Supplementary Tables

**Supplementary Table 2.** Number of brown bear occurrences (n = 3,226,206) and percentage by each type of data/source and group. We include the identifier of the research group (ID).

| ID | Tracks and Observations | GPS telemetry | VHF telemetry | Genetic and hairtraps | Cameras | Mixed |
|---|---|---|---|---|---|---|
| 1 | 13103 | 0 | 0 | 0 | 0 | 0 |
| 2 | 0 | 88617 | 0 | 0 | 0 | 0 |
| 3 | 0 | 13106 | 0 | 356 | 0 | 0 |
| 4 | 0 | 109414 | 0 | 0 | 0 | 0 |
| 5 | 0 | 0 | 0 | 1598 | 0 | 0 |
| 6 | 0 | 0 | 0 | 0 | 0 | 375 |
| 7 | 2654 | 38222 | 0 | 346 | 0 | 0 |
| 8 | 0 | 0 | 0 | 22247 | 0 | 0 |
| 9 | 0 | 0 | 0 | 0 | 0 | 1150 |
| 10 | 3918 | 108672 | 0 | 334 | 0 | 0 |
| 11 | 556 | 3737 | 473 | 0 | 0 | 0 |
| 12 | 2234 | 5583 | 5584 | 0 | 0 | 0 |
| 13 | 0 | 29933 | 0 | 0 | 0 | 0 |
| 14 | 47 | 0 | 0 | 0 | 0 | 0 |
| 15 | 1859 | 115397 | 0 | 0 | 0 | 0 |
| 16 | 201 | 0 | 0 | 0 | 130 | 0 |
| 17 | 103 | 0 | 0 | 0 | 0 | 0 |
| 18 | 114 | 0 | 0 | 0 | 0 | 0 |
| 19 | 0 | 1959497 | 0 | 0 | 0 | 0 |
| 20 | 0 | 0 | 0 | 0 | 0 | 2094 |
| 21 | 2318 | 0 | 0 | 0 | 0 | 0 |
| 22 | 0 | 1309 | 0 | 0 | 0 | 0 |
| 23 | 0 | 183133 | 0 | 0 | 0 | 0 |
| 24 | 1736 | 0 | 10165 | 0 | 0 | 0 |
| 25 | 0 | 165165 | 0 | 0 | 0 | 0 |
| 26 | 0 | 0 | 700 | 0 | 0 | 0 |
| 27 | 270 | 36728 | 0 | 0 | 0 | 0 |
| 28 | 6 | 1438 | 0 | 0 | 91 | 0 |
| 29 | 2861 | 124561 | 0 | 0 | 0 | 0 |
| 30 | 640 | 0 | 0 | 0 | 0 | 0 |
| 31 | 91 | 0 | 0 | 0 | 0 | 0 |
| 32 | 2408 | 0 | 0 | 0 | 0 | 0 |
| 33 | 401 | 0 | 0 | 0 | 0 | 0 |
| 34 | 0 | 0 | 0 | 0 | 106 | 0 |
| 35 | 598 | 0 | 0 | 0 | 0 | 0 |
| 36 | 0 | 159827 | 0 | 0 | 0 | 0 |
| TOTAL | 36118 | 3144339 | 16922 | 24881 | 327 | 1151 |
| Percentage | 1.12 | 97.46 | 0.52 | 0.77 | 0.01 | 0.11 |

**Supplementary Table 3.** Number of pixels of 1x1 km with presence of brown bear by subpopulation (n presences all systems), the number of pixels in terrestrial systems (n presences in terrestrial), the number of presences used to fit the brown bear models at habitat scale (including models comparing biotic variables and brown bear model at habitat scale; n presences selected to train the habitat models), the number of presences used to validate the brown bear models at habitat scale (brown bear model at habitat scale; n presences selected to validate the habitat model).

| Subpopulation | n presences all systems | n presences in terrestrial | n presences selected to train the habitat models | n presences selected to validate the habitat model |
|---|---|---|---|---|
| *Alpine* | 3091 | 3087 | 1600 | 400 |
| *Baltic* | 2449 | 2447 | 1600 | 400 |
| *Cantabrian* | 2194 | 2186 | 1600 | 400 |
| *Eastern Carpathian* | 8208 | 8202 | 1600 | 400 |
| *Western Carpathian* | 3191 | 3190 | 1600 | 400 |
| *Caucasian* | 2132 | 2132 | 1600 | 400 |
| *Apennine* | 1151 | 1148 | 918 | 230 |
| *East Balkan* | 2278 | 2278 | 1600 | 400 |
| *Pindus* | 3612 | 3603 | 1600 | 400 |
| *Dinaric* | 8954 | 8947 | 1600 | 400 |
| *Karelian* | 18490 | 1793 | 1600 | 400 |
| *Pyrenees* | 1289 | 1289 | 1031 | 258 |
| *Scandinavian* | 45119 | 44151 | 1600 | 400 |
| *Turkey* | 471 | 471 | 377 | 94 |

**Supplementary Table 10.** Sum of rates of Estimated Dietary Energy Content (*rEDEC*) described at the species level for each subpopulation ($rEDEC_{Subp}$).

| Subpopulation | $rEDEC_{Subp}$ |
|---|---|
| *Alpine* | 0.33 |
| *Baltic* | 0.75 |
| *Cantabrian* | 0.96 |
| *Eastern Carpathian* | 0.77 |
| *Western Carpathian* | 0.80 |
| *Caucasian* | 0.20 |
| *Apennine* | 0.83 |
| *East Balkan* | 1.00 |
| *Pindus* | 0.51 |
| *Dinaric* | 0.81 |
| *Karelian* | 0.90 |
| *Pyrenees* | 0.50 |
| *Scandinavian* | 0.86 |
| *Turkey* | 0.63 |

**Supplementary Table 11.** Sum of rates of Estimated Dietary Energy Content, *rEDEC*, described at the species level for each subpopulation by diet group.

| Subpopulation | Invertebr. | Reprod. plant material | Unknown plant material and others | Veget. plant material | Vertebr. |
|---|---|---|---|---|---|
| *Apennine* | 0.29 | 87.46 | 0.00 | 1.49 | 10.75 |
| *Cantabrian* | 1.30 | 69.49 | 8.73 | 2.60 | 17.80 |
| *Scandinavian* | 3.35 | 39.97 | 0.34 | 5.18 | 51.16 |
| *Alpine* | 16.85 | 23.75 | 21.36 | 5.05 | 32.99 |
| *Eastern Carpathian* | 0.00 | 87.73 | 0.00 | 0.71 | 11.56 |
| *East Balkan* | 1.50 | 94.12 | 0.00 | 0.00 | 4.37 |
| *Dinaric* | 0.00 | 91.76 | 0.00 | 0.00 | 8.24 |
| *Turkey* | 0.00 | 17.55 | 67.18 | 0.00 | 15.27 |
| *Pyrenees* | 0.00 | 51.65 | 5.87 | 4.04 | 38.44 |
| *Pindos* | 0.00 | 86.86 | 0.00 | 1.25 | 11.89 |
| *Baltic* | 0.53 | 88.42 | 0.04 | 0.38 | 10.63 |
| *Karelian* | 6.37 | 56.69 | 0.00 | 0.00 | 36.94 |
| *Western Carpathian* | 0.60 | 80.10 | 0.00 | 0.00 | 19.29 |
| *Caucasian* | 0.00 | 32.15 | 0.00 | 0.00 | 67.85 |

**Supplementary Table 12.** Rates (from the total described at the species level for each subpopulation) of Estimated Dietary Energy Content, *rEDEC*, by origin (Wild/Human) for each subpopulation.

| **Subpopulation** | **Wild** | **Human** |
|---|---|---|
| *Apennine* | 76.58 | 21.67 |
| *Cantabrian* | 83.20 | 16.70 |
| *Scandinavian* | 61.66 | 38.34 |
| *Alpine* | 85.05 | 14.95 |
| *Eastern Carpathian* | 47.48 | 52.52 |
| *East Balkan* | 7.02 | 92.98 |
| *Dinaric* | 43.57 | 56.43 |
| *Turkey* | 94.40 | 5.60 |
| *Pyrenees* | 79.89 | 20.11 |
| *Pindos* | 59.21 | 40.72 |
| *Baltic* | 41.03 | 58.97 |
| *Karelian* | 97.64 | 2.36 |
| *Western Carpathian* | 57.23 | 42.77 |
| *Caucasian* | 49.24 | 50.76 |

**Supplementary Table 34.** Statistics, minimum, median, mean and maximum of the evaluations of species distribution models for wild food species.

| | TSS | Sensitivity | Specificity |
|---|---|---|---|
| Minimum | 0.11 | 36.23 | 44.33 |
| Median | 0.47 | 77.88 | 69.37 |
| Mean | 0.47 | 78.04 | 69.43 |
| Maximum | 0.74 | 97.47 | 91.27 |

**Supplementary Table 36.** Average importance for the variables used to fit species distribution models for wild food species.

| Variable | Importance |
|---|---|
| *Clim_1* | 0.32 |
| *Clim_12* | 0.12 |
| *Clim_15* | 0.12 |
| *Clim_7* | 0.24 |
| *LC_2* | 0.25 |
| *LC_3* | 0.04 |
| *LC_4* | 0.14 |
| *LC_5* | 0.03 |
| *LC_7* | 0.01 |
| *LC_8* | 0.00 |
| *LC_9* | 0.00 |

**Supplementary Table 38.** Mean habitat suitability change (in percentage) for each food category and scenario.

| Scenario | Diet category | Habitat suitability change (%) |
|---|---|---|
| SSP1-2.6 | reproductive_plant_material | -42.87 |
| SSP1-2.6 | unknown_plant_material_and_others | -31.83 |
| SSP1-2.6 | vegetative_plant_material | -32.44 |
| SSP1-2.6 | invertebrates | 18.44 |
| SSP1-2.6 | vertebrates | -46.94 |
| SSP3-6.0 | reproductive_plant_material | -45.72 |
| SSP3-6.0 | unknown_plant_material_and_others | -34.20 |
| SSP3-6.0 | vegetative_plant_material | -36.32 |
| SSP3-6.0 | invertebrates | 25.77 |
| SSP3-6.0 | vertebrates | -44.65 |
| SSP5-8.5 | reproductive_plant_material | -62.08 |
| SSP5-8.5 | unknown_plant_material_and_others | -53.49 |
| SSP5-8.5 | vegetative_plant_material | -38.03 |
| SSP5-8.5 | invertebrates | 8.21 |
| SSP5-8.5 | vertebrates | -52.06 |

**Supplementary Table 40.** Performance for univariable species distribution models (frequentist binomial GLMMs) fitted for each food category explaining the distribution of brown bear. For each food category (All species, Reproductive plants, Vegetative plants, Unknown plants, Invertebrate and Vertebrates), we fitted two models, one with *Biotic variables* and other using *Biotic_binary variables*. To select variables representing the biotic interaction for the brown bear model at habitat scale we excluded the group including all species as it was highly correlated with other biotic variables. We used 19,926 presences and 19,926 absences to fit these models (Supplementary Table 3). We show the Akaike information criterion (AIC; a lower AIC value indicates a better fit of the model) for each model, and for each diet category, we show the difference in AIC between both univariable models (AIC *Biotic_binary variables* – AIC *Biotic variables*; positive values in the difference indicate that *Biotic variables* models have a better fit than *Biotic_binary variables*) and the difference of the AIC *Biotic variables* with the best univariable model considering biotic information (Delta AIC *Biotic variables*). In addition we calculated a *Null* model (a GLMM with only the interaction term) as a reference and we report the AIC and delta of this model.

| **Group of species** | **Variable name** | | **AIC models** | | **AIC *Biotic_binary variables* – AIC *Biotic variables*** | **Delta AIC *Biotic variables*** |
|---|---|---|---|---|---|---|
| | ***Biotic variables*** | ***Biotic_binary variables*** | ***Biotic variables*** | ***Biotic_binary variables*** | | |
| *All species* | $Bio_{All\_species}$ | $Bio\_binary_{All\_species}$ | 54594.33 | 55070.34 | 476.01 | 0.00 |
| *Reprod plant* | $Bio_{Reprod\_plant}$ | $Bio\_binary_{Reprod\_plant}$ | 54604.92 | 54959.81 | 354.89 | 10.59 |
| *Veget plant* | $Bio_{Unknown\_plant}$ | $Bio\_binary_{Unknown\_plant}$ | 55238.03 | 55180.96 | -57.08 | 643.70 |
| *Unknown plant* | $Bio_{Unknown\_plant}$ | $Bio\_binary_{Unknown\_plant}$ | 55029.78 | 55141.39 | 111.61 | 435.45 |
| *Invertebrates* | $Bio_{Invertebrates}$ | $Bio\_binary_{Invertebrates}$ | 55199.62 | 55252.57 | 52.95 | 605.28 |
| *Vertebrates* | $Bio_{Vertebrates}$ | $Bio\_binary_{Vertebrates}$ | 55165.40 | 55227.99 | 62.59 | 571.07 |
| *Null* | - | - | 55250.60 | | - | 656.27 |

**Supplementary Table 42.** Fitted bayesian models (BMs) explaining brown bear distribution. We show the name for each model (Model name), whether abiotic and/or biotic variables are included in the models (Predictors included) and the type of model, which indicate if it is a simple Bayesian model (with uninformative priors, BSM) or a hierarchical Bayesian model (where climatic variables are using priors based in the model of brown bear at range scale, BHM). In addition to the three BMs described in the main text: (1) a model with abiotic and biotic predictors, (2) a model with abiotic predictors only, and (3) a model with biotic predictors only, we calculated two other alternative models in order to evaluate the effect of combining different data. Those alternative models (shaded in grey) were alternatives to both BHMs, the model with abiotic and biotic predictors ($SHM_{ABI}$), and the model with abiotic predictors only ($SHM_{AI}$), the $SHM_{ABC}$ and $SHM_{AC}$ respectively which used the same predictors but were BSMs only using data from brown bear *Occurrence Database* (Methods).

| **Model name** | **Predictors included** | **Type of model** |
|---|---|---|
| $SHM_{ABI}$ | Abiotic and biotic | BHM |
| $SHM_{ABC}$ | Abiotic and biotic | BSM |
| $SHM_{AI}$ | Abiotic | BHM |
| $SHM_{AC}$ | Abiotic | BSM |
| $SHM_{BC}$ | Biotic | BSM |
| $SHM_{Null}$ | Only the intercept | BSM |

**Supplementary Table 43.** Performance for bayesian models (BMs) explaining brown bear distribution. For each model we show the model name (See Supplementary Table 42), estimates, standard error (SE) the information criterion WAIC (which is just -2 * elpd_waic, i.e., converted to deviance scale), the expected log pointwise predictive density (elpd_waic), the effective number of parameters (p_waic), and for the mean of the sample average posterior predictive distribution of the outcome (mean_PPD); and delta WAIC values. A plausible mean indicates when compared with the mean (y; $y_{mean} = 0.4708$) does not mean that it is a good model, but a not plausible mean indicates a wrong model. Diagnostics for Pareto smoothed importance sampling (PSIS) indicated that all pareto k estimates were good ($k < 0.5$). We used 19,926 presences and 19,926 absences to fit these models (Supplementary Table 3). Alternative models are shaded in grey (See Supplementary Table 42).

| | WAIC | | | elpd_WAIC | | p_WAIC | | mean_PPD | |
|---|---|---|---|---|---|---|---|---|---|
| **Model name** | **Estimate** | **SE** | **Delta WAIC** | **Estimate** | **SE** | **Estimate** | **SE** | **Estimate** | **SD** |
| $SHM_{ABI}$ | 52710.9 | 92.8 | 0 | -26355.5 | 46.4 | 26.6 | 0.3 | 0.50 | 0.003 |
| $SHM_{ABC}$ | 52787.2 | 95.2 | 76.3 | -26393.6 | 47.6 | 28.1 | 0.3 | 0.50 | 0.004 |
| $SHM_{AI}$ | 53048.0 | 84.7 | 337.1 | -26524.0 | 42.3 | 22.5 | 0.3 | 0.50 | 0.003 |
| $SHM_{AC}$ | 53119.1 | 88.6 | 408.2 | -26559.6 | 44.3 | 24.3 | 0.3 | 0.50 | 0.003 |
| $SHM_{BC}$ | 54322.1 | 60.3 | 1611.2 | -27161.0 | 30.2 | 18.0 | 0.1 | 0.50 | 0.003 |
| $SHM_{Null}$ | 55250.6 | 0.0 | 2539.7 | -27625.3 | 0.0 | 2.0 | 0.0 | 0.50 | 0.004 |

**Supplementary Table 44.** Results for the Bayesian hierarchical model using abiotic and biotic factors to explain brown bear distribution ($SHM_{ABI}$; see Supplementary Table 42). We report model coefficients (best estimates and their SE), Monte Carlo standard error (MCSE), confident intervals (10%, 50% and 90%), number of effective sample size (Neff) and the potential scale reduction factor on split chains (Rhat; at convergence Rhat=1).

| | **mean** | **mcse** | **sd** | **10%** | **50%** | **90%** | **Neff** | **Rhat** |
|---|---|---|---|---|---|---|---|---|
| Intercept | -15.51 | 0.02 | 0.919 | -16.72 | -15.49 | -14.35 | 3619 | 1.000 |
| *Clim_3* | 18.85 | 0.03 | 1.655 | 16.70 | 18.89 | 20.95 | 3786 | 0.999 |
| *Clim_4* | 6.01 | 0.01 | 0.504 | 5.36 | 6.01 | 6.65 | 3838 | 1.000 |
| *Clim_8* | 0.96 | 0.00 | 0.053 | 0.89 | 0.96 | 1.03 | 3527 | 1.001 |
| *Clim_10* | 3.34 | 0.00 | 0.172 | 3.12 | 3.34 | 3.56 | 3207 | 1.000 |
| *Clim_3_c* | -38.03 | 0.05 | 2.984 | -41.82 | -38.05 | -34.23 | 3725 | 0.999 |
| *Clim_4_c* | -3.18 | 0.00 | 0.275 | -3.53 | -3.18 | -2.84 | 3808 | 1.000 |
| *Clim_8_c* | -0.50 | 0.00 | 0.030 | -0.54 | -0.50 | -0.46 | 3661 | 1.000 |
| *Clim_10_c* | -1.16 | 0.00 | 0.051 | -1.22 | -1.16 | -1.09 | 3118 | 1.000 |
| *LC_1* | -0.06 | 0.00 | 0.016 | -0.08 | -0.06 | -0.04 | 6188 | 0.999 |
| *LC_3* | 0.20 | 0.00 | 0.017 | 0.18 | 0.20 | 0.22 | 3701 | 0.999 |
| *LC_4* | 0.28 | 0.00 | 0.019 | 0.25 | 0.28 | 0.30 | 3836 | 1.000 |
| *Nat. Landscape* | 0.09 | 0.00 | 0.019 | 0.07 | 0.09 | 0.11 | 3949 | 1.000 |
| $Bio_{Reprod\_plant}$ | 0.18 | 0.00 | 0.016 | 0.15 | 0.18 | 0.20 | 5758 | 1.000 |
| $Bio_{Invertebrates}$ | -0.14 | 0.00 | 0.017 | -0.17 | -0.14 | -0.12 | 6011 | 0.999 |
| $Bio_{Unknown\_plant}$ | 0.22 | 0.00 | 0.038 | 0.17 | 0.22 | 0.27 | 4489 | 1.000 |
| $Bio_{Vertebrates}$ | 0.09 | 0.00 | 0.014 | 0.08 | 0.09 | 0.11 | 5715 | 1.000 |

**Supplementary Table 45.** Results for the simple Bayesian model (no hierarchical) using abiotic and biotic factors to explain brown bear distribution (*$SHM_{ABC}$*; see Supplementary Table 42). We report model coefficients (best estimates and their SE), Monte Carlo standard error (MCSE), confident intervals (10%, 50% and 90%), number of effective sample size (Neff) and the potential scale reduction factor on split chains (Rhat; at convergence Rhat=1).

| | **mean** | **mcse** | **sd** | **10%** | **50%** | **90%** | **Neff** | **Rhat** |
|---|---|---|---|---|---|---|---|---|
| Intercept | -6.28 | 0.02 | 0.768 | -7.26 | -6.28 | -5.30 | 2498 | 1.001 |
| *Clim_3* | 4.17 | 0.02 | 1.327 | 2.45 | 4.18 | 5.86 | 3155 | 1.001 |
| *Clim_4* | 5.54 | 0.03 | 1.562 | 3.55 | 5.54 | 7.56 | 2312 | 1.000 |
| *Clim_8* | 1.17 | 0.00 | 0.071 | 1.08 | 1.17 | 1.26 | 3743 | 1.000 |
| *Clim_10* | 4.42 | 0.01 | 0.358 | 3.96 | 4.42 | 4.89 | 2952 | 1.001 |
| *Clim_3_c* | -9.04 | 0.04 | 2.185 | -11.87 | -8.99 | -6.32 | 3536 | 1.000 |
| *Clim_4_c* | -3.10 | 0.02 | 0.908 | -4.27 | -3.10 | -1.94 | 2411 | 0.999 |
| *Clim_8_c* | -0.62 | 0.00 | 0.040 | -0.67 | -0.62 | -0.57 | 3662 | 1.000 |
| *Clim_10_c* | -1.47 | 0.00 | 0.107 | -1.60 | -1.47 | -1.33 | 2959 | 1.000 |
| *LC_1* | -0.06 | 0.00 | 0.016 | -0.08 | -0.06 | -0.04 | 5557 | 1.000 |
| *LC_3* | 0.19 | 0.00 | 0.016 | 0.17 | 0.19 | 0.21 | 4279 | 1.000 |
| *LC_4* | 0.26 | 0.00 | 0.018 | 0.24 | 0.26 | 0.28 | 3755 | 1.001 |
| *Nat. Landscape* | 0.10 | 0.00 | 0.018 | 0.07 | 0.10 | 0.12 | 4202 | 1.000 |
| $Bio_{Reprod\_plant}$ | 0.17 | 0.00 | 0.017 | 0.15 | 0.17 | 0.19 | 5231 | 1.000 |
| $Bio_{Invertebrates}$ | -0.15 | 0.00 | 0.017 | -0.17 | -0.15 | -0.13 | 5790 | 1.000 |
| $Bio_{Unknown\_plant}$ | 0.18 | 0.00 | 0.038 | 0.14 | 0.18 | 0.23 | 3451 | 1.000 |
| $Bio_{Vertebrates}$ | 0.10 | 0.00 | 0.014 | 0.08 | 0.10 | 0.12 | 5635 | 1.000 |

**Supplementary Table 46.** Results for the Bayesian hierarchical model using abiotic factors to explain brown bear distribution (*$SHM_{AI}$*; see Supplementary Table 42). We report model coefficients (best estimates and their SE), Monte Carlo standard error (MCSE), confident intervals (10%, 50% and 90%), number of effective sample size (Neff) and the potential scale reduction factor on split chains (Rhat; at convergence Rhat=1).

| | **mean** | **mcse** | **sd** | **10%** | **50%** | **90%** | **Neff** | **Rhat** |
|---|---|---|---|---|---|---|---|---|
| Intercept | -14.93 | 0.02 | 0.936 | -16.09 | -14.94 | -13.71 | 3347 | 1.002 |
| *Clim_3* | 17.89 | 0.03 | 1.656 | 15.76 | 17.87 | 20.02 | 3301 | 1.000 |
| *Clim_4* | 5.83 | 0.01 | 0.493 | 5.20 | 5.84 | 6.48 | 3803 | 1.000 |
| *Clim_8* | 0.93 | 0.00 | 0.052 | 0.87 | 0.93 | 1.00 | 3827 | 1.000 |
| *Clim_10* | 3.51 | 0.00 | 0.168 | 3.30 | 3.51 | 3.72 | 3564 | 1.001 |
| *Clim_3_c* | -39.67 | 0.05 | 3.040 | -43.57 | -39.66 | -35.76 | 3536 | 0.999 |
| *Clim_4_c* | -3.41 | 0.00 | 0.267 | -3.75 | -3.41 | -3.05 | 3711 | 0.999 |
| *Clim_8_c* | -0.49 | 0.00 | 0.029 | -0.53 | -0.50 | -0.46 | 3597 | 1.000 |
| *Clim_10_c* | -1.23 | 0.00 | 0.050 | -1.30 | -1.23 | -1.17 | 3515 | 1.000 |
| *LC_1* | -0.06 | 0.00 | 0.016 | -0.08 | -0.06 | -0.05 | 5598 | 1.000 |
| *LC_3* | 0.23 | 0.00 | 0.016 | 0.21 | 0.23 | 0.25 | 4345 | 0.999 |
| *LC_4* | 0.29 | 0.00 | 0.018 | 0.27 | 0.29 | 0.32 | 4239 | 0.999 |
| *Nat. Landscape* | 0.12 | 0.00 | 0.019 | 0.10 | 0.12 | 0.15 | 4316 | 0.999 |

**Supplementary Table 47.** Results for the simple Bayesian model using abiotic factors to explain brown bear distribution ($SHM_{AC}$; see Supplementary Table 42). We report model coefficients (best estimates and their SE), Monte Carlo standard error (MCSE), confident intervals (10%, 50% and 90%), number of effective sample size (Neff) and the potential scale reduction factor on split chains (Rhat; at convergence Rhat=1).

| | **mean** | **mcse** | **sd** | **10%** | **50%** | **90%** | **Neff** | **Rhat** |
|---|---|---|---|---|---|---|---|---|
| Intercept | -6.33 | 0.01 | 0.735 | -7.26 | -6.33 | -5.38 | 2878 | 1.000 |
| *Clim_3* | 3.22 | 0.02 | 1.363 | 1.50 | 3.23 | 4.98 | 3159 | 1.000 |
| *Clim_4* | 5.72 | 0.03 | 1.560 | 3.74 | 5.72 | 7.72 | 2126 | 1.000 |
| *Clim_8* | 1.13 | 0.00 | 0.072 | 1.04 | 1.13 | 1.22 | 3337 | 1.000 |
| *Clim_10* | 5.12 | 0.01 | 0.345 | 4.68 | 5.12 | 5.56 | 3122 | 1.000 |
| *Clim_3_c* | -9.40 | 0.04 | 2.175 | -12.24 | -9.41 | -6.62 | 3745 | 0.999 |
| *Clim_4_c* | -3.64 | 0.02 | 0.904 | -4.81 | -3.62 | -2.49 | 2214 | 1.000 |
| *Clim_8_c* | -0.61 | 0.00 | 0.041 | -0.67 | -0.61 | -0.56 | 3252 | 1.000 |
| *Clim_10_c* | -1.70 | 0.00 | 0.103 | -1.83 | -1.69 | -1.56 | 3204 | 1.000 |
| *LC_1* | -0.06 | 0.00 | 0.016 | -0.09 | -0.06 | -0.04 | 5232 | 0.999 |
| *LC_3* | 0.22 | 0.00 | 0.016 | 0.20 | 0.22 | 0.24 | 3898 | 1.000 |
| *LC_4* | 0.27 | 0.00 | 0.018 | 0.24 | 0.27 | 0.29 | 3551 | 0.999 |
| *Nat. Landscape* | 0.13 | 0.00 | 0.019 | 0.10 | 0.13 | 0.15 | 3732 | 1.000 |

**Supplementary Table 48.** Results for the simple Bayesian model using biotic factors to explain brown bear distribution ($SHM_{BC}$; see Supplementary Table 42). We report model coefficients (best estimates and their SE), Monte Carlo standard error (MCSE), confident intervals (10%, 50% and 90%), number of effective sample size (Neff) and the potential scale reduction factor on split chains (Rhat; at convergence Rhat=1).

| | **mean** | **mcse** | **sd** | **10%** | **50%** | **90%** | **Neff** | **Rhat** |
|---|---|---|---|---|---|---|---|---|
| Intercept | 0.02 | 0.01 | 0.164 | -0.19 | 0.02 | 0.23 | 569 | 1.007 |
| $Bio_{Reprod\_plant}$ | 0.33 | 0.00 | 0.015 | 0.31 | 0.33 | 0.35 | 3580 | 1.000 |
| $Bio_{Invertebrates}$ | -0.19 | 0.00 | 0.016 | -0.21 | -0.19 | -0.16 | 3572 | 1.000 |
| $Bio_{Unknown\_plant}$ | 0.34 | 0.00 | 0.039 | 0.29 | 0.34 | 0.40 | 2554 | 1.000 |
| $Bio_{Vertebrates}$ | 0.05 | 0.00 | 0.014 | 0.03 | 0.05 | 0.07 | 3237 | 1.001 |

**Supplementary Table 54.** Results for the validation of the best Bayesian model (BM) explaining brown bear distribution, the Bayesian hierarchical model using abiotic and biotic factors (*$SHM_{ABI}$*; see Supplementary Table 42). to validate the model. We show the values to correctly classify the pseudo-absences of brown bear (true negative rate; TNR), to correctly classify the presences of brown bear (true positive rate; TPR) and classification accuracy (Acc.) at European scale and by subpopulation. We used an independent subset of data from the *Ocurrence Database*, the validation subset, which contains 4,982 presences and 4,982 absences to validate these models (Supplementary Table 3).

| | **Europe** | **Alpine** | **Baltic** | **Cantabrian** | **Eastern Carpathian** | **Western Carpathian** | **Caucasian** | **Apennine** | **East Balkan** | **Pindus** | **Dinaric** | **Karelian** | **Pyrenees** | **Scandinavian** | **Turkey** |
|---|---|---|---|---|---|---|---|---|---|---|---|---|---|---|---|
| **TNR** | 0.25 | 0.29 | 0.27 | 0.26 | 0.33 | 0.15 | 0.38 | 0.08 | 0.28 | 0.29 | 0.25 | 0.05 | 0.44 | 0.17 | 0.45 |
| **TPR** | 0.87 | 0.92 | 0.89 | 0.92 | 0.78 | 0.95 | 0.75 | 0.97 | 0.85 | 0.74 | 0.94 | 0.98 | 0.76 | 0.91 | 0.74 |
| **Acc.** | 0.56 | 0.61 | 0.58 | 0.59 | 0.56 | 0.55 | 0.56 | 0.53 | 0.56 | 0.52 | 0.59 | 0.52 | 0.60 | 0.54 | 0.60 |

**Supplementary Table 59.** Name of climatic variables and definition.

| **Name** | **Definition** |
|---|---|
| *Clim_1* | Annual mean temperature [1/10°C] |
| *Clim_2* | Mean diurnal range [1/10°C] |
| *Clim_3* | Isothermality * 100 |
| *Clim_4* | Temperature seasonality * 100 |
| *Clim_5* | Maximum Temperature of warmest month [1/10°C] |
| *Clim_6* | Minimum Temperature of coldest month [1/10°C] |
| *Clim_7* | Temperature Annual Range [1/10°C] |
| *Clim_8* | Mean Temperature of wettest quarter [1/10°C] |
| *Clim_9* | Mean Temperature of driest quarter [1/10°C] |
| *Clim_10* | Mean Temperature of warmest quarter [1/10°C] |
| *Clim_11* | Mean Temperature of coldest quarter [1/10°C] |
| *Clim_12* | Annual precipitation amount [mm] |
| *Clim_13* | Precipitation of wettest month [mm] |
| *Clim_14* | Precipitation of driest month [mm] |
| *Clim_15* | Precipitation Seasonality [coefficient of variation * 100] |
| *Clim_16* | Precipitation of wettest quarter [mm] |
| *Clim_17* | Precipitation of driest quarter [mm] |
| *Clim_18* | Precipitation of warmest quarter [mm] |
| *Clim_19* | Precipitation of coldest quarter [mm] |

**Supplementary Table 60.** Name of land cover variables, definition and original categories.

| Name | Definition | ID CCI-LC classification included |
|---|---|---|
| *LC_1* | Percentage of urban areas | 1, 190 |
| *LC_2* | Percentage of cultivated and pasture areas | 2, 230,231, 232, 3 |
| *LC_3* | Percentage of broadleaved forest | 50, 60, 61, 62 |
| *LC_4* | Percentage of needle-leaved forest | 70, 71, 72, 80, 82, 90 |
| *LC_5* | Percentage of Shrubland | 100, 110, 120, 121, 122 |
| *LC_7* | Sparse vegetation | 150, 152, 153 |
| *LC_8* | Percentage of rangeland areas | 4 |
| *LC_9* | Percentage of bare areas | 200, 201, 202 |
| *Nat. Landscape* | Percentage of natural areas at landscape scale. Value obtained from the application of a gaussian smoothing kernel function moving window of 11 x 11 km to a layer of percentage of natural areas. | 50, 60, 61, 62, 70, 71, 72, 80, 81, 82, 90, 100, 110, 120, 121, 122, 130, 140, 150, 152, 153, 160, 170, 180, 200, 201, 202 |

**Supplementary Table 61.** Evaluation of univariable models for no correlated bioclimatic variables at range scale using historical distribution and historical climate. We indicate with a grey shadow the best four models based on Akaike information criterion (AIC). These variables were used to model the habitat of brown bear in all models including climatic variables.

| Model/Variable | AIC |
|---|---|
| *Clim_2* | 38237.81 |
| *Clim_3* | 27249.12 |
| *Clim_4* | 27296.31 |
| *Clim_8* | 29483.18 |
| *Clim_10* | 21682.73 |
| *Clim_13* | 30705.14 |
| *Clim_14* | 36276.78 |
| *Clim_15* | 37577.52 |
| *Clim_18* | 33587.15 |
| *Clim_19* | 37659.84 |

**Supplementary Table 62.** Results for the brown bear species distribution model for the historical range, based on bayesian GLM predicting presence (using historical distribution in Eurasia) as a function of bioclimatic variables. We report model coefficients (best estimates and their SE), Monte Carlo standard error (MCSE), confident intervals (10%, 50% and 90%), number of effective sample size (Neff) and the potential scale reduction factor on split chains (Rhat; at convergece Rhat=1).

| | **mean** | **mcse** | **sd** | **10%** | **50%** | **90%** | **Neff** | **Rhat** |
|---|---|---|---|---|---|---|---|---|
| Intercept | -9.55 | 0.01 | 0.772 | -10.53 | -9.55 | -8.56 | 9631.00 | 1.00 |
| *Clim_3* | 18.10 | 0.03 | 2.854 | 14.49 | 18.07 | 21.75 | 8364.00 | 1.00 |
| *Clim_4* | 7.52 | 0.01 | 0.814 | 6.49 | 7.50 | 8.57 | 7706.00 | 1.00 |
| *Clim_8* | 0.62 | 0.00 | 0.121 | 0.46 | 0.62 | 0.78 | 9577.00 | 1.00 |
| *Clim_10* | 2.95 | 0.00 | 0.268 | 2.61 | 2.94 | 3.30 | 7431.00 | 1.00 |
| *Clim_3_c* | -17.76 | 0.04 | 3.902 | -22.81 | -17.68 | -12.85 | 7933.00 | 1.00 |
| *Clim_4_c* | -2.57 | 0.00 | 0.329 | -3.00 | -2.56 | -2.15 | 7501.00 | 1.00 |
| *Clim_8_c* | -0.38 | 0.00 | 0.057 | -0.45 | -0.37 | -0.30 | 9753.00 | 1.00 |
| *Clim_10_c* | -1.10 | 0.00 | 0.085 | -1.21 | -1.09 | -0.99 | 7544.00 | 1.00 |

**Supplementary Table 63.** Sample average posterior predictive distribution of the outcome for the for the brown bear species distribution model for the historical range. A plausible mean indicates when compared with the mean (y; $y_{mean}$ = 0.3794) does not mean that it is a good model, but a not plausible mean indicates a wrong model.

| | **mean** | **sd** | **10%** | **50%** | **90%** |
|---|---|---|---|---|---|
| mean_PPD | 0.38 | 0.013 | 0.36 | 0.38 | 0.40 |

**Supplementary Table 64.** Estimates and standard error (SE) for the expected log pointwise predictive density (elpd_waic), the effective number of parameters (p_waic) and the information criterion waic (which is just -2 * elpd_waic, i.e., converted to deviance scale) for the brown bear species distribution model for the historical range. Diagnostics for Pareto smoothed importance sampling (PSIS) indicated that all pareto k estimates were good (k < 0.5).

| | **Estimate** | **SE** |
|---|---|---|
| **elpd_WAIC** | -1030.0 | 19.7 |
| **p_WAIC** | 8.0 | 0.3 |
| **WAIC** | 2060.0 | 39.5 |

**Supplementary Table 65.** Estimates and standard error (SE) for the expected log pointwise predictive density (elpd_waic), the effective number of parameters (p_waic) and the information criterion waic (which is just -2 * elpd_waic, i.e., converted to deviance scale) for a null species distribution model at range scale, based on bayesian GLM predicting the distribution with only the intercept.

| | **Estimate** | **SE** |
|---|---|---|
| **elpd_WAIC** | -1388.3 | 10.9 |
| **p_WAIC** | 1.0 | 0.0 |
| **WAIC** | 2776.5 | 21.8 |

**Supplementary Table 67.** Correlation among land use variables with current data of brown bear.

| | ***LC_1*** | ***LC_2*** | ***LC_3*** | ***LC_4*** | ***LC_5*** | ***LC_8*** | ***LC_9*** | ***Nat. Landscape*** |
|---|---|---|---|---|---|---|---|---|
| ***LC_1*** | 1.00 | | | | | | | |
| ***LC_2*** | 0.05 | 1.00 | | | | | | |
| ***LC_3*** | -0.04 | -0.27 | 1.00 | | | | | |
| ***LC_4*** | -0.05 | -0.37 | -0.39 | 1.00 | | | | |
| ***LC_5*** | -0.02 | -0.19 | -0.13 | -0.11 | 1.00 | | | |
| ***LC_8*** | -0.01 | 0.00 | -0.12 | -0.18 | -0.08 | 1.00 | | |
| ***LC_9*** | -0.01 | -0.05 | -0.07 | -0.08 | -0.01 | -0.03 | 1.00 | |
| ***Nat. Landscape*** | -0.08 | **-0.71** | 0.24 | 0.57 | 0.13 | -0.30 | 0.01 | 1.00 |

**Supplementary Table 68.** Evaluation of univariable models for land use variables at using the *Ocurrence Database*. We indicate with a grey shadow the best four models based in Akaike information criterion (AIC). These variables were used to model the habitat of brown bear in all models including land use variables.

| Variable | AIC | |
|---|---|---|
| *LC_1* | 55156.95 | |
| *LC_2* | 54887.07 | Correlated with *Nat. Landscape* |
| *LC_3* | 55019.35 | |
| *LC_4* | 54640.32 | |
| *LC_5* | 55240.58 | |
| *LC_8* | 55223.67 | |
| *LC_9* | 55147.85 | |
| *Nat. Landscape* | 54342.63 | |
| *Null* | 55250.60 | |

### Supplementary Figures

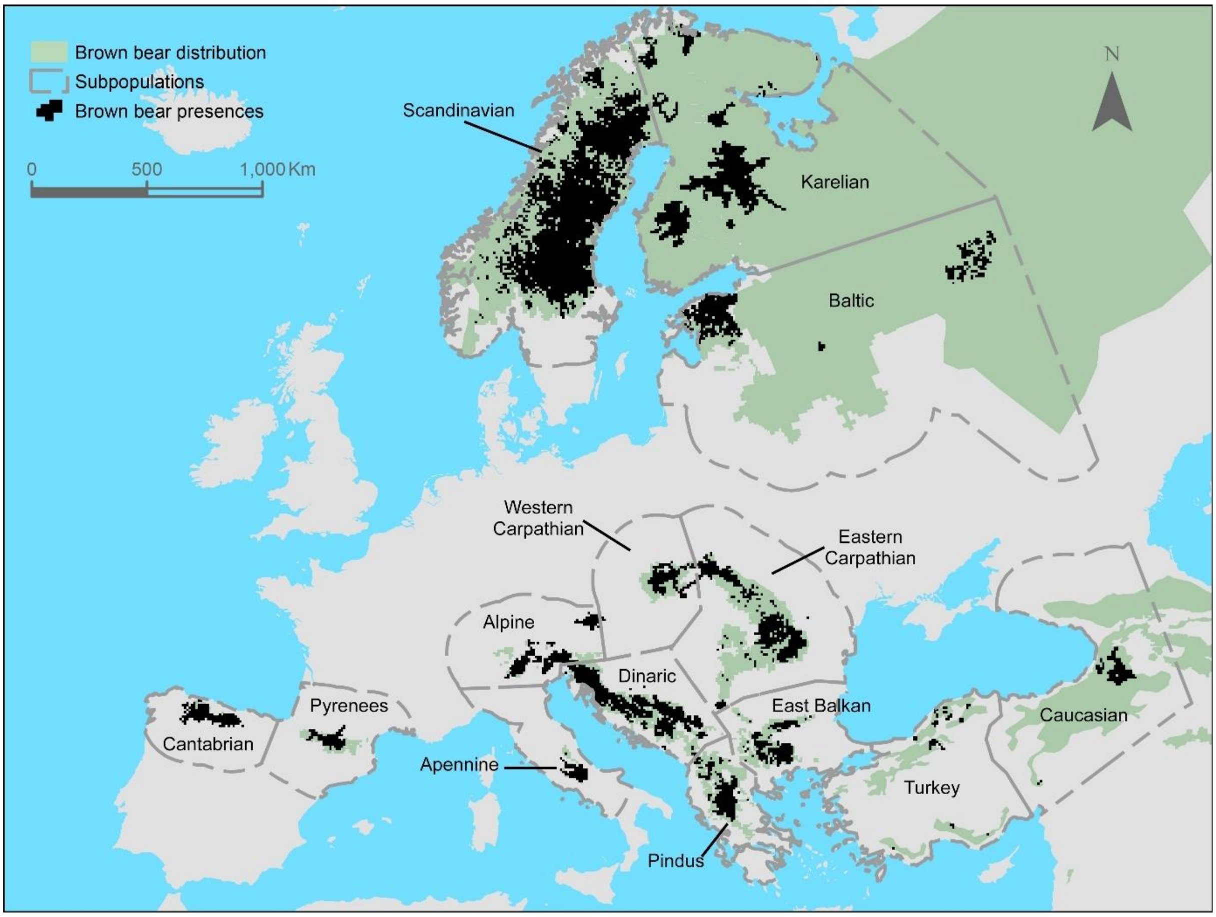


**Supplementary Figure 1.** Map showing 10×10 km cells with presence of brown bear from the *Occurrence Database* which includes brown bear presences at high resolution (1×1km).

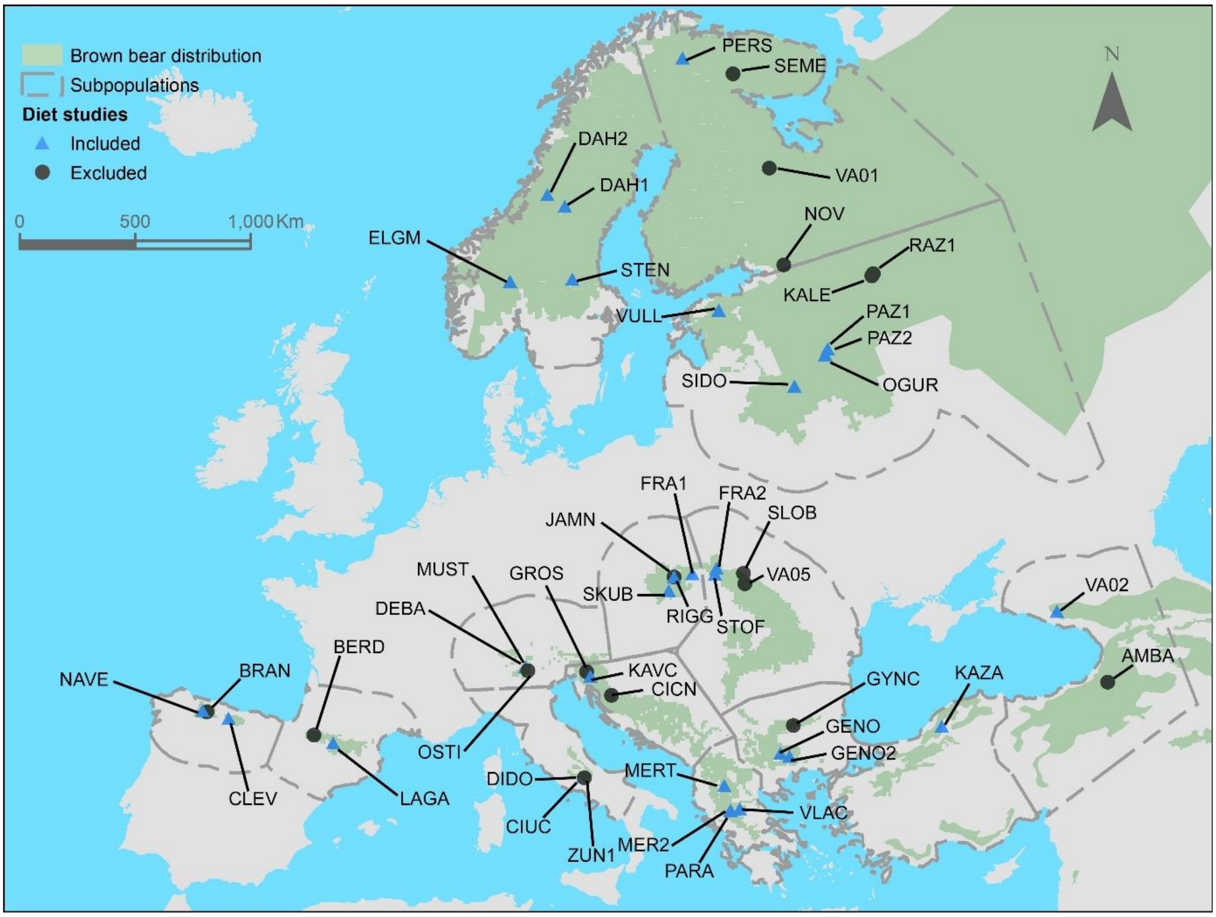


**Supplementary Figure 2.** Map showing the location of 47 studies of brown bear diet found in the review. For consistency with climate and land use data, we selected 31 studies conducted between 1989 and 2018 which had sufficient taxonomic resolution (genus and/or species; Supplementary Table 4).

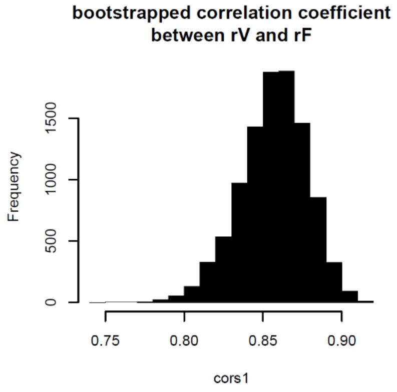


**Supplementary Figure 3**. Bootstrapped correlation between relative frequency and relative volume.

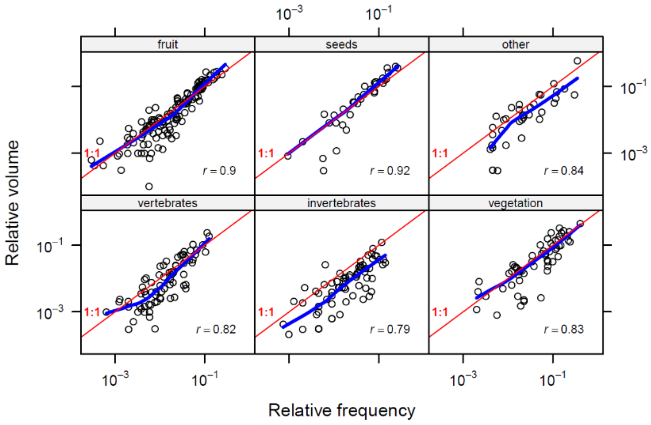


**Supplementary Figure 4.** Empirical relationships between relative frequency and relative volume.

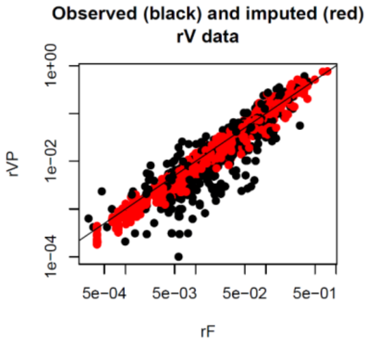


**Supplementary Figure 5.** Relationships between relative volume with relative frequency of observed and imputed data.

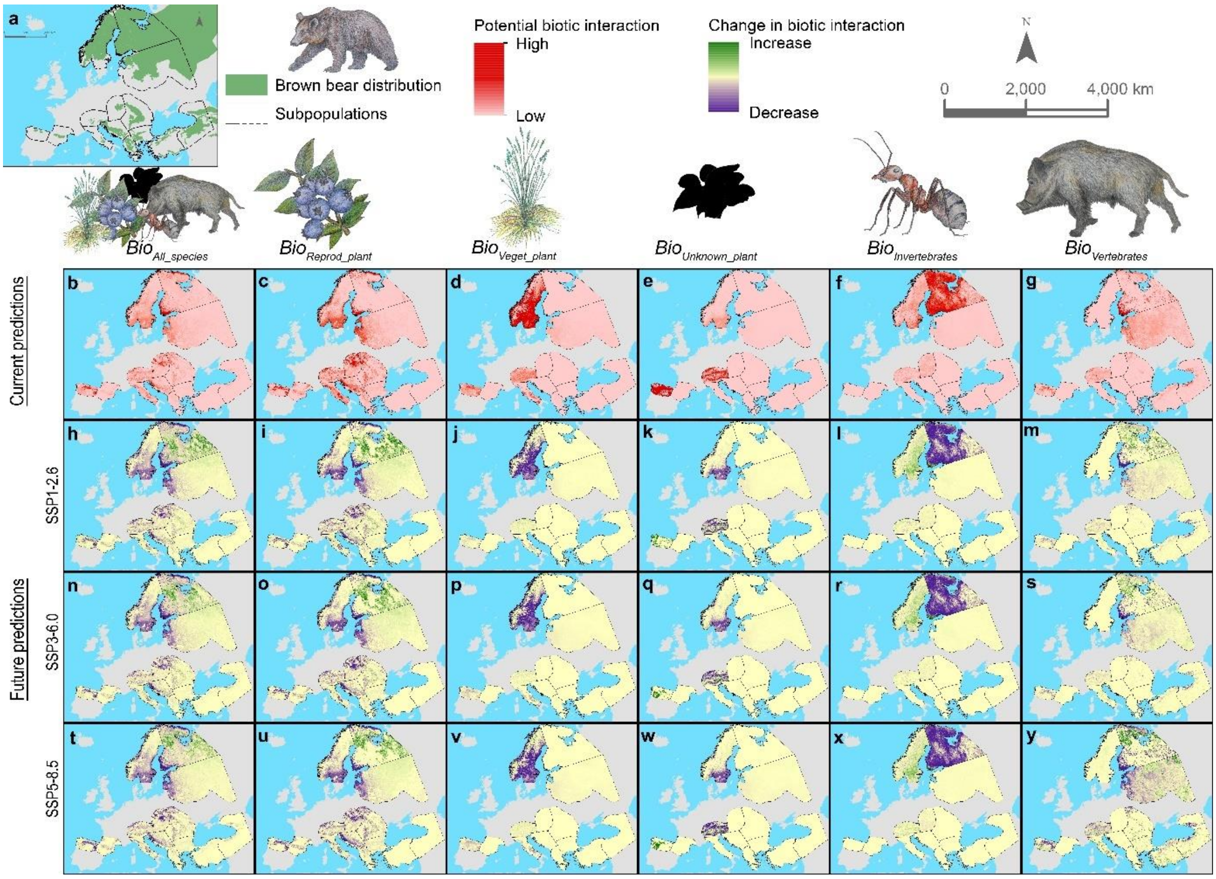


**Supplementary Figure 6.** Map showing the current biotic variables and the change in biotic variables for the future SSPs. **a** Current distribution of the brown bear in Europe. **b-g** Current prediction for *Biotic variables*. **h-y** Future prediction for *Biotic variables* for the three shared socioeconomic pathways considered.

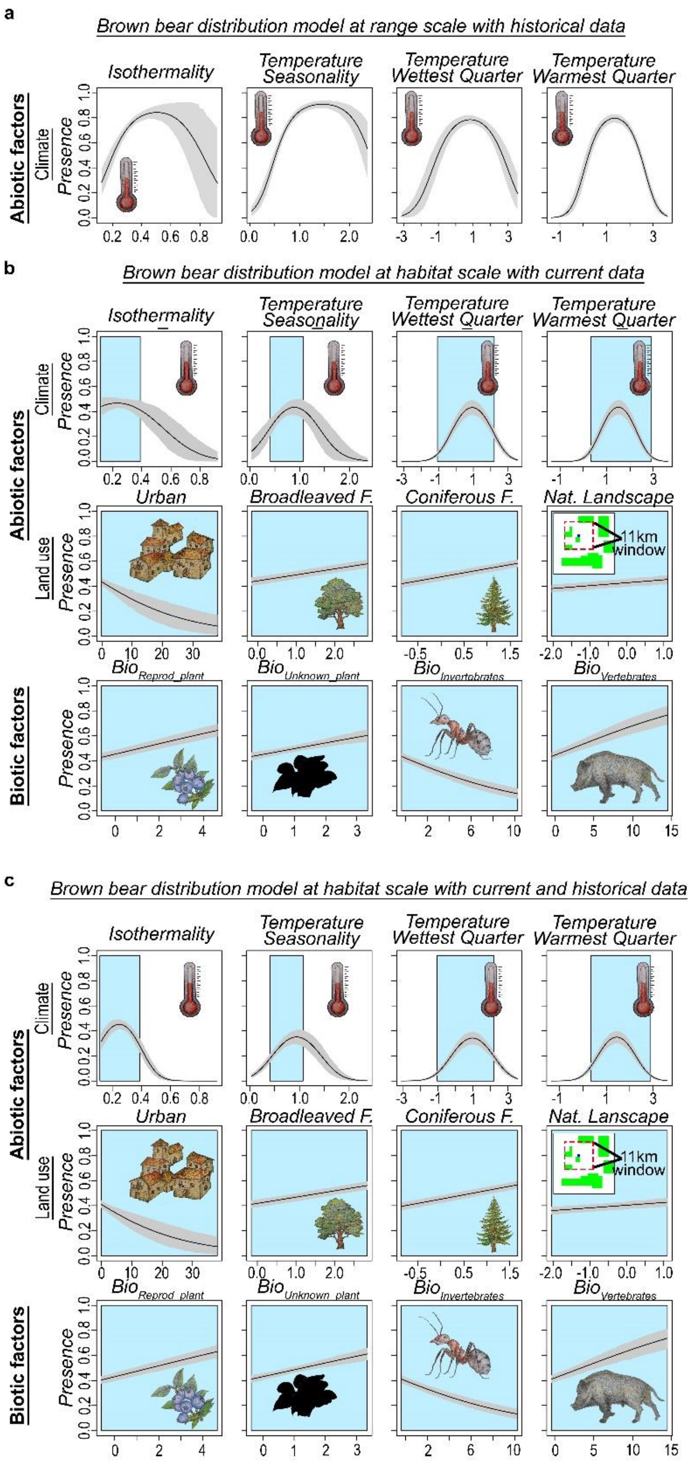


**Supplementary Figure 7.** Response plot of the three Bayesian models explaining the distribution of the brown bear. **a** Bayesian model at range scale using the historical distribution of brown bear and historical climate variables as predictors. **b** Simple Bayesian model (no hierarchical) using abiotic and biotic factors to explain brown bear distribution. **c** Distribution model for brown bear including both abiotic and biotic factors was fitted combining both historical (*Range Database*) and current data (*Occurrence Database*). The continuous line represents the mean response value, and the grey area shows the model uncertainty (95% confidence interval). The blue area indicates the range of values of the current data. Isothermality (*Clim_3*), temperature seasonality (*Clim_4*), mean temperature of the wettest quarter (*Clim_5*) and mean temperature of the warmest quarter (*Clim_10*).

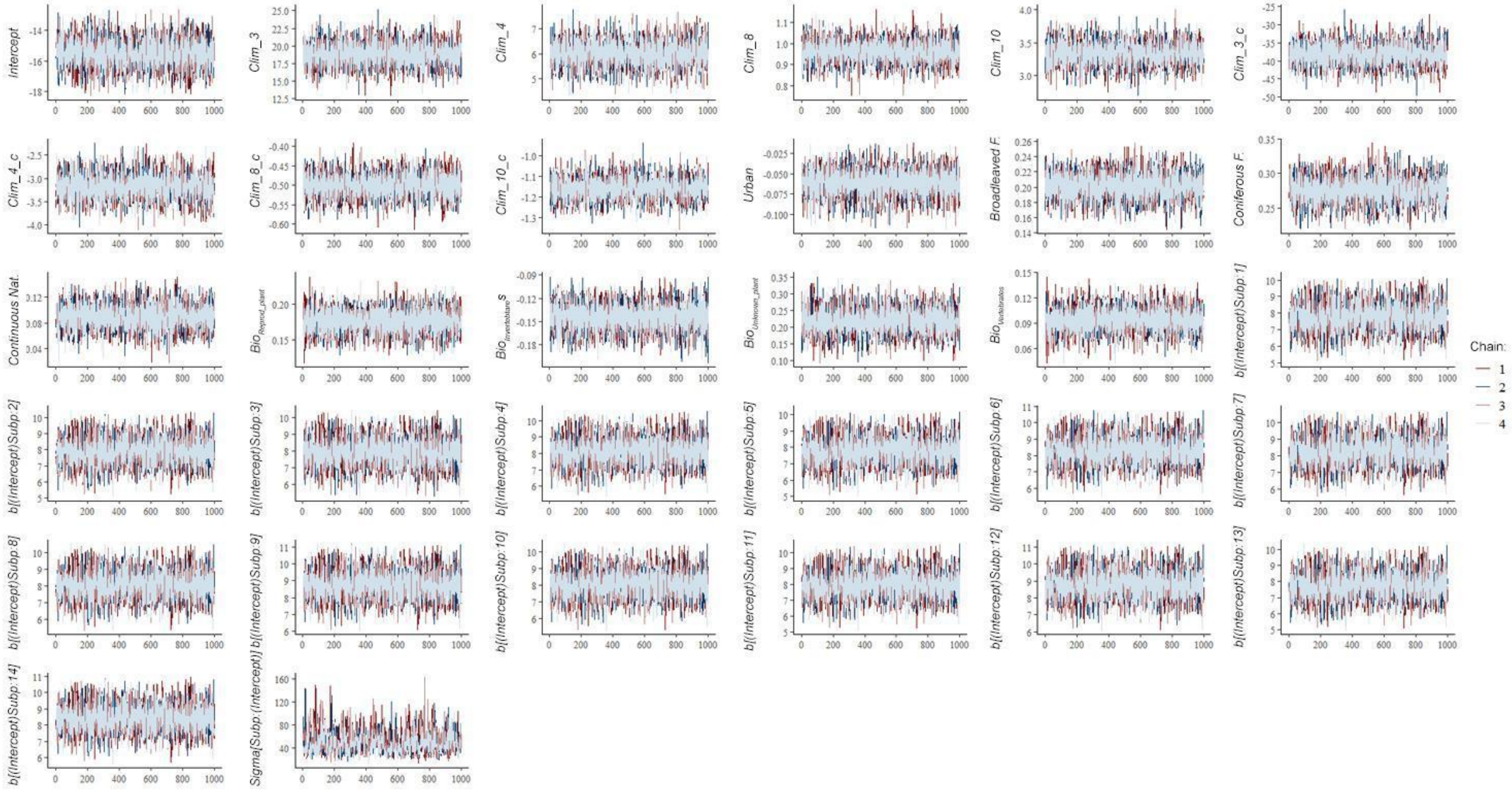


**Supplementary Figure 8.** Chains for the Bayesian model of brown bear habitat with abiotic and biotic factors combining data.

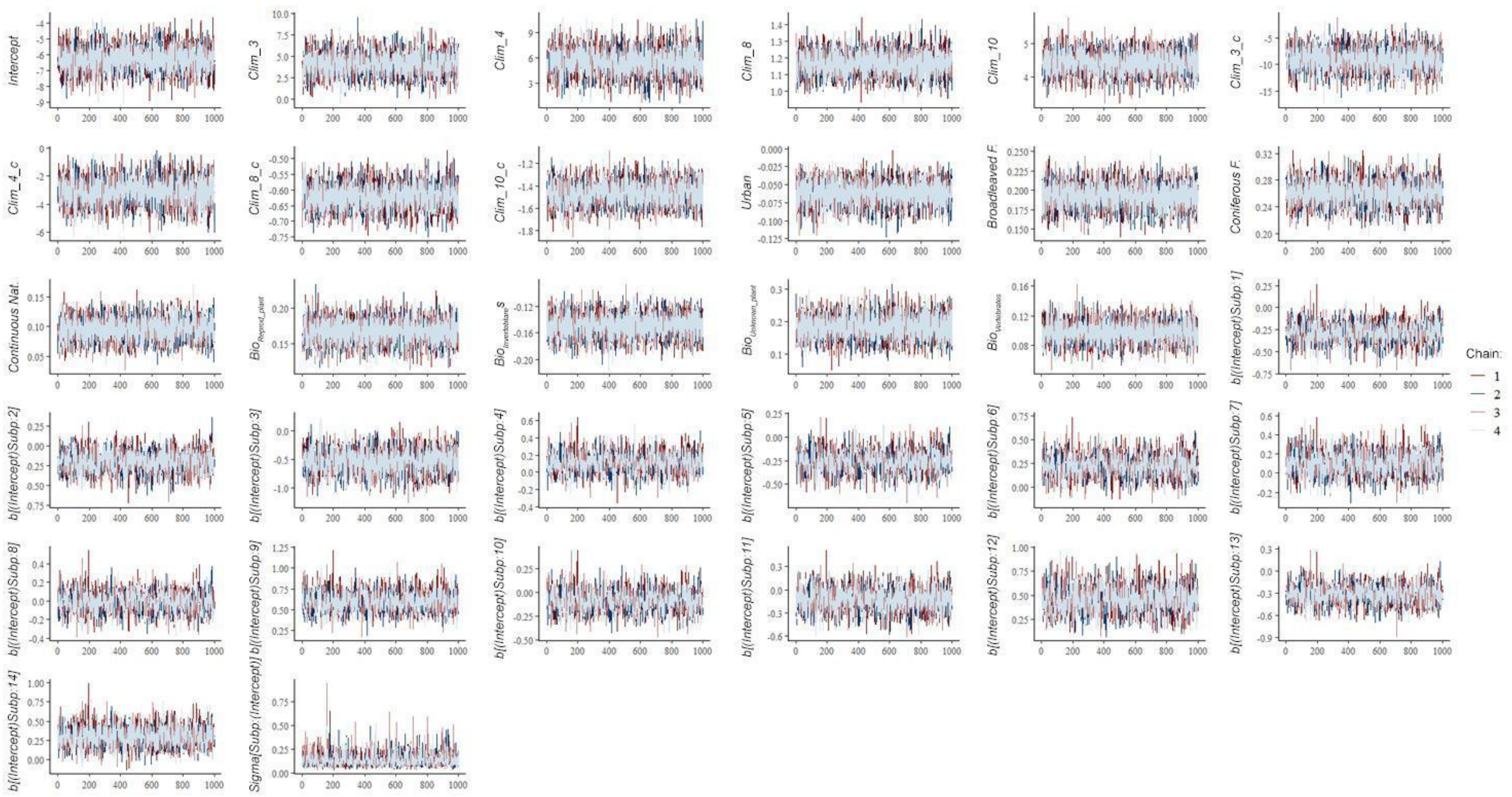


**Supplementary Figure 9.** Chains for the Bayesian model of brown bear habitat with abiotic and biotic factors using current data.

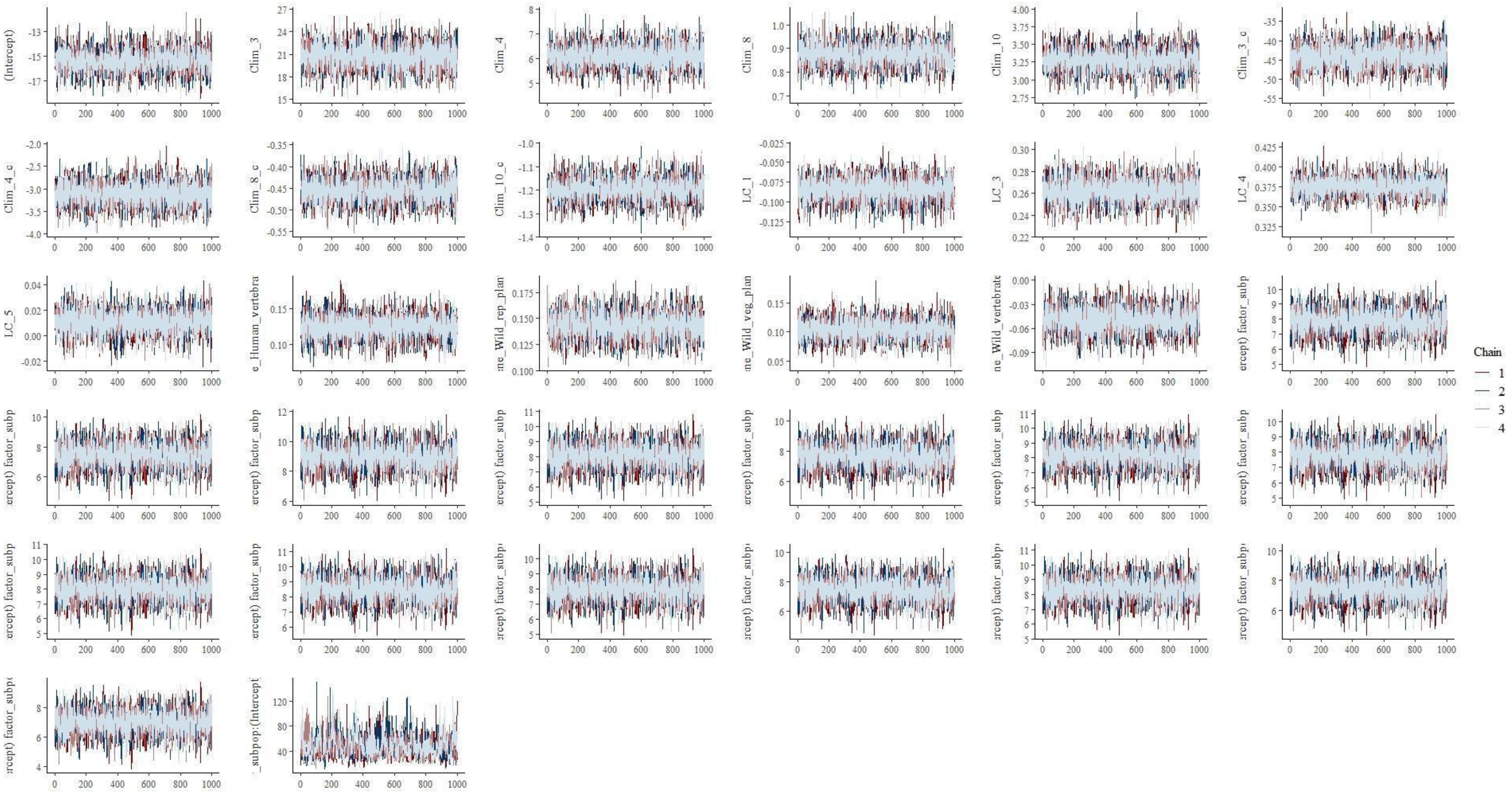

**Supplementary Figure 10.** Chains for the Bayesian model of brown bear habitat with abiotic factors combining data.

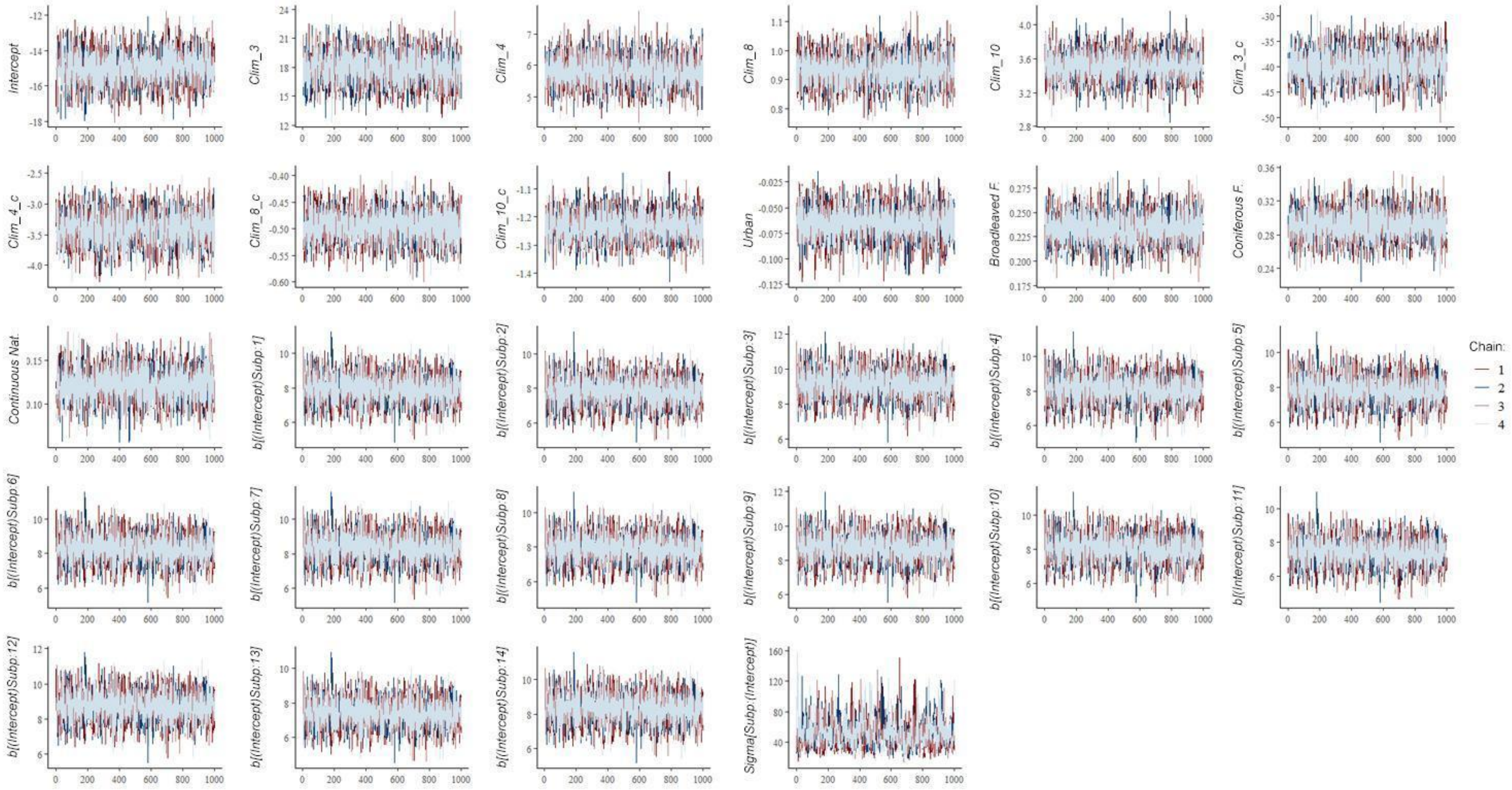


**Supplementary Figure 11.** Chains for the Bayesian model of brown bear habitat with abiotic factors using current data.

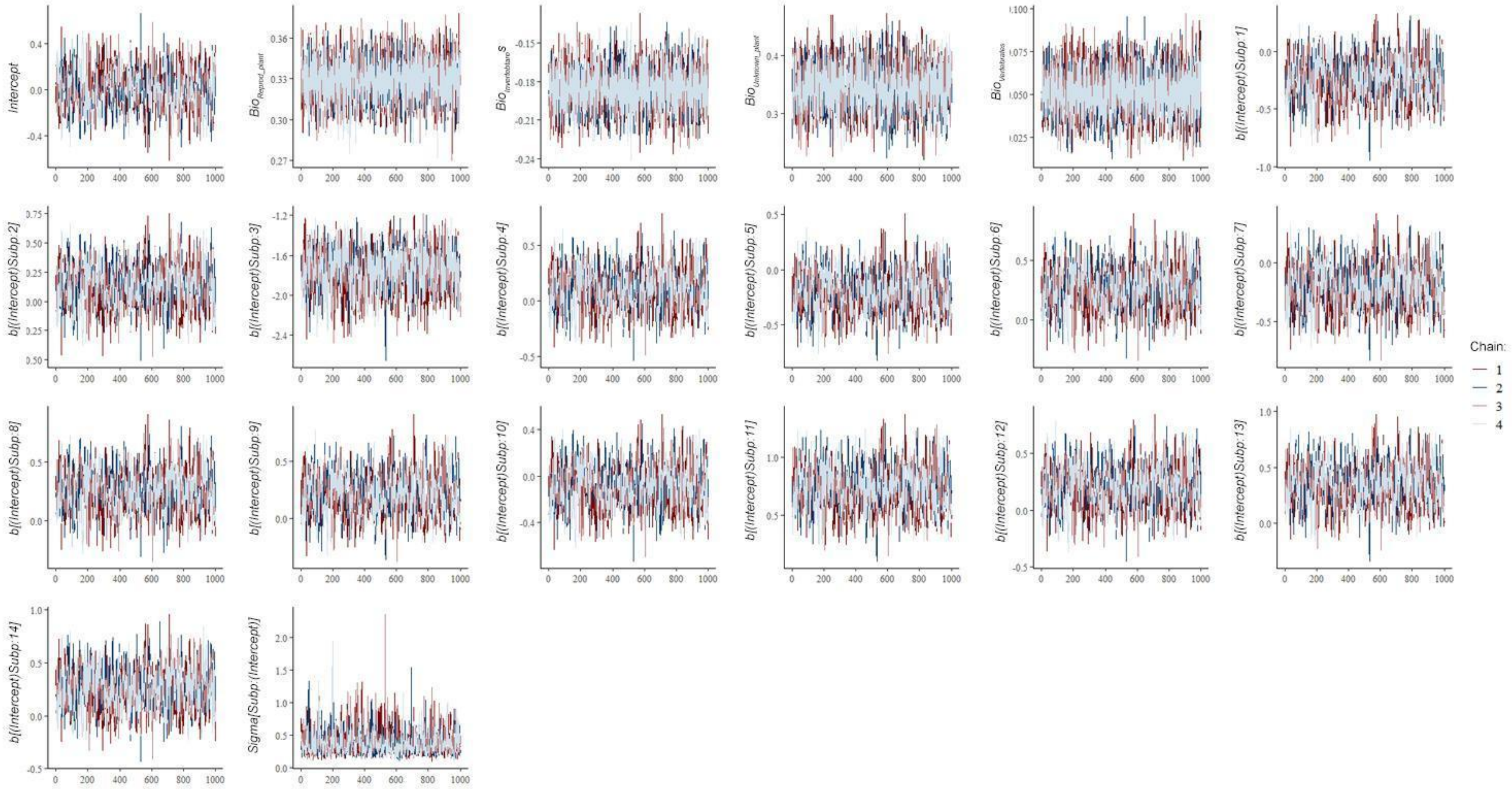


**Supplementary Figure 12.** Chains for the Bayesian model of brown bear habitat with biotic factors.

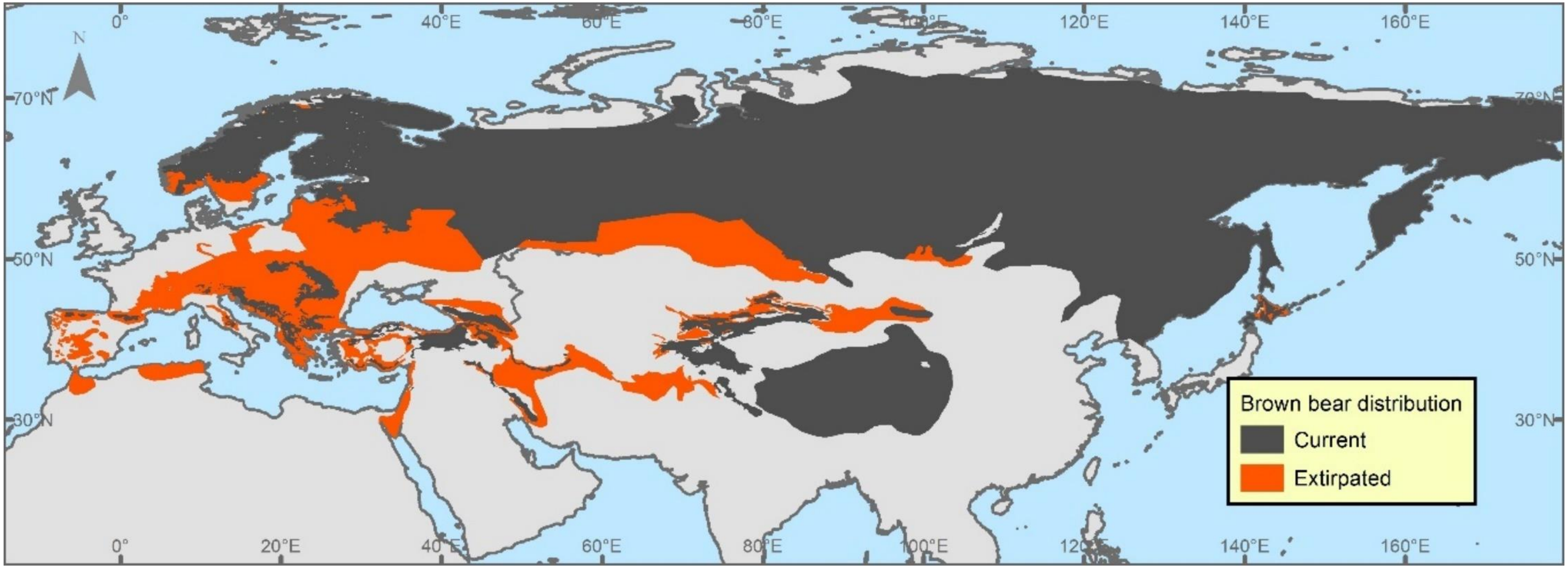


**Supplementary Figure 13.** Historical distribution (Current and Extirpated areas) of brown bear in Eurasia used in the Species Distribution Model range. Current distribution was obtained from the IUCN Red List spatial data and extirpated distribution was build based on IUCN data in its majority and completed with other sources (See methods section).

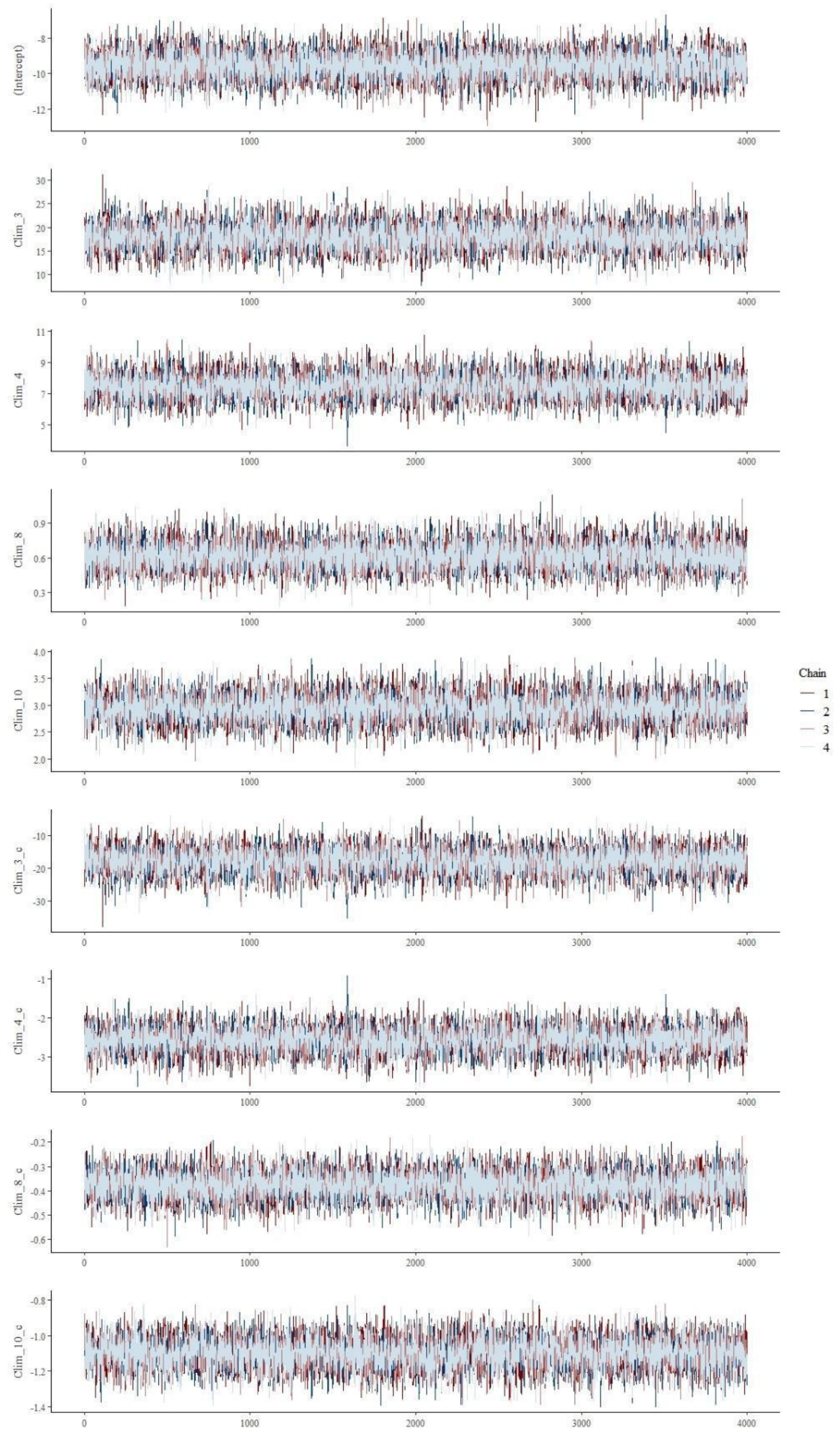


**Supplementary Figure 14.** Chains for the Species Distribution Model at range scale, the $SDM_{Range}$

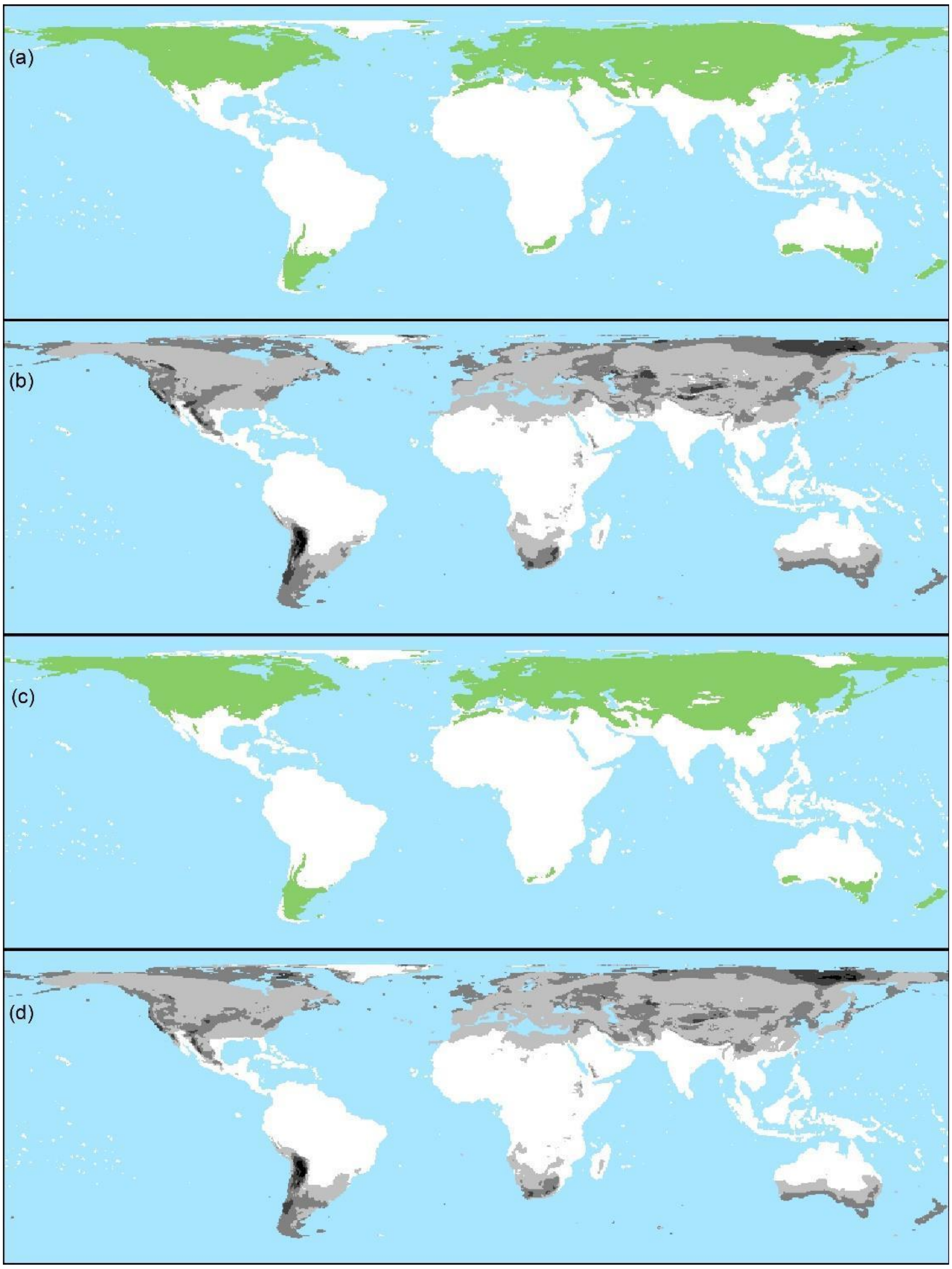


**Supplementary Figure 15.** Maps for the Bayesian model at range scale explaining brown bear distribution. We show the predicted distribution (a), and uncertainty in mean predicted (b) for historical climate data. And the predicted distribution (c), and uncertainty in mean predicted (d) for current climate data.

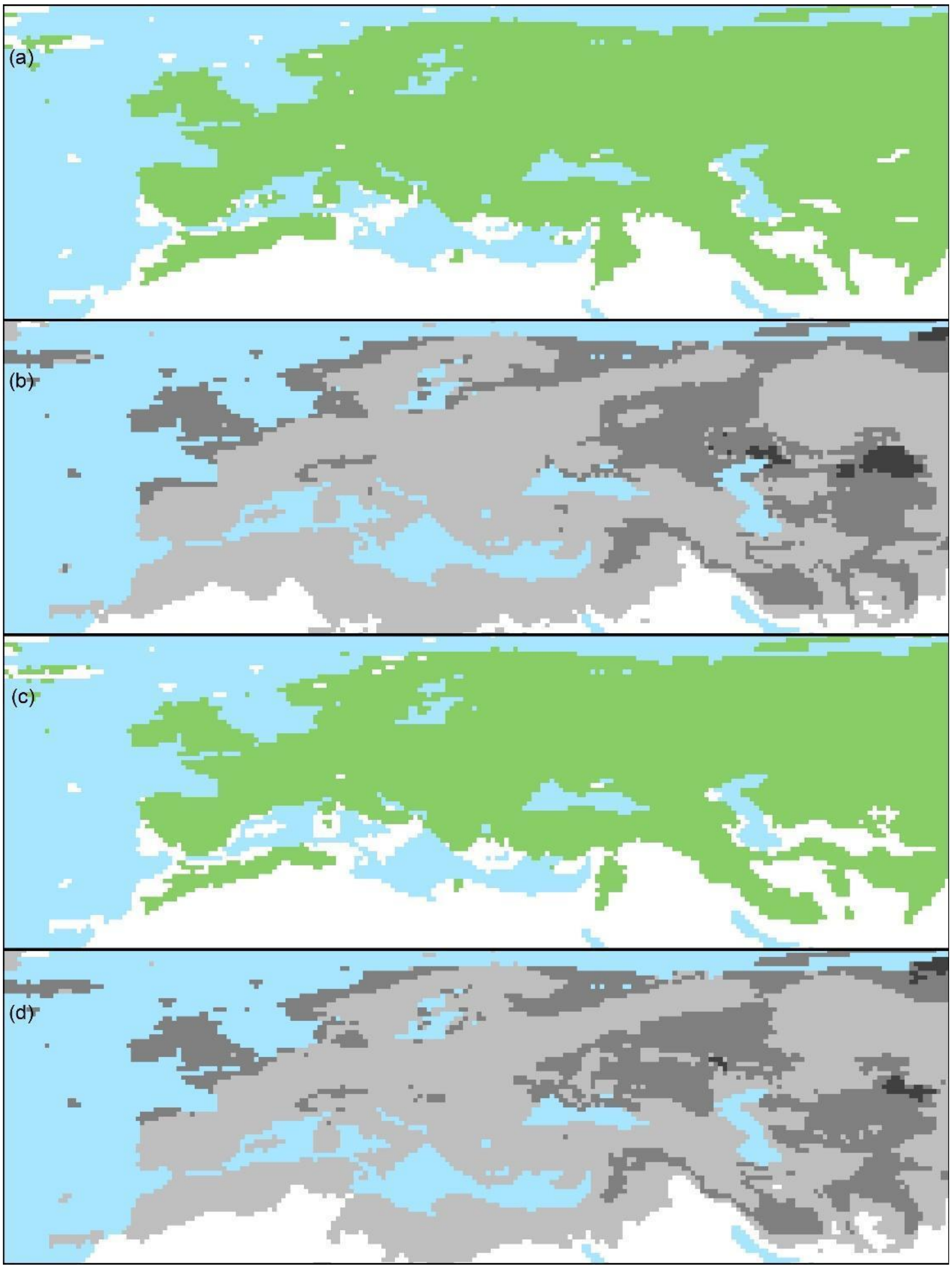


**Supplementary Figure 16.** Maps for the Bayesian model at range scale explaining brown bear distribution. We show the predicted distribution (a), and uncertainty in mean predicted (b) for historical climate data. And the predicted distribution (c), and uncertainty in mean predicted (d) for current climate data.

109 IPCC. *Climate Change 2013: The physical science basis. Contribution of Working Group I to the Fifth Assessment Report of the Intergovermental Panel on Climate Change*. (Cambridge University Press, 2014).

110 Organization, W. M. <http://www.wmo.int/pages/index_en.html> (

111 Hijmans, R. J., Phillips, S., Leathwick , J. & Elith, J. (2020).
